# eIF5B and eIF1A remodel human translation initiation complexes to mediate ribosomal subunit joining

**DOI:** 10.1101/2021.12.09.471821

**Authors:** Christopher P. Lapointe, Rosslyn Grosely, Masaaki Sokabe, Carlos Alvarado, Jinfan Wang, Elizabeth Montabana, Nancy Villa, Byung-Sik Shin, Thomas E. Dever, Christopher S. Fraser, Israel S. Fernández, Joseph D. Puglisi

## Abstract

Joining of the ribosomal subunits at a translation start site on a messenger RNA during initiation commits the ribosome to synthesize a protein. Here, we combined single-molecule spectroscopy and structural methods using an *in vitro* reconstituted system to examine how the human ribosomal subunits join. Single-molecule fluorescence revealed when universally-conserved eukaryotic initiation factors (eIFs) eIF1A and eIF5B associate with and depart from initiation complexes. Guided by single-molecule dynamics, we examined initiation complexes that contained both eIF1A and eIF5B using single-particle electron cryo-microscopy. The resulting structure illuminated how eukaryote-specific contacts between eIF1A and eIF5B remodel the initiation complex to orient initiator tRNA in a conformation compatible with ribosomal subunit joining. Collectively, our findings provide a quantitative and architectural framework for the molecular choreography orchestrated by eIF1A and eIF5B during human translation initiation.

## INTRODUCTION

Human protein synthesis initiates when ribosomal subunits identify a translation start site on messenger RNA (mRNA) and assemble a functional 80S ribosome. The multi-step process has been outlined biochemically and is mediated by numerous eukaryotic initiation factors (eIFs) and the initiator aminoacyl-tRNA (Met-tRNA_i_^Met^) (*1-3*). To begin, the 43S pre-initiation complex (43S PIC) – the small (40S) ribosomal subunit bound by eIFs 1, 1A, 3, 5, and the eIF2–GTP–Met-tRNA_i_^Met^ complex – loads onto the mRNA, which is facilitated by eIF4 proteins bound near the m^7^G-cap (*4, 5*). Directional 5’ to 3’ movement of the now 48S initiation complex along the 5’ untranslated region scans for translation start sites in favorable sequence contexts (*6, 7*). A start codon (typically AUG) correctly placed at the ribosomal P site triggers remodeling of the complex from an ‘open’ scanning-permissive conformation to a ‘closed’ scanning-arrested one. The transition involves repositioning of the 40S head and Met-tRNA_i_^Met^, ejection of eIF1 and the eIF2-GDP complex, and relocation of eIF5 (*7, 8*). To complete initiation, a GTPase, eIF5B, catalyzes joining of the large (60S) ribosomal subunit to form the 80S ribosome (*9-11*). Subsequent GTP hydrolysis-dependent departure of eIF5B enables protein synthesis to commence.

What allows ribosomal subunits to join remains unclear. One challenge is that the acceptor stem of Met-tRNA_i_^Met^ must be slotted into the correct narrow channel on the incoming 60S subunit. Proper Met-tRNA_i_^Met^ orientation avoids steric clashes and poises its aminoacylated 3’ end for accommodation in the peptidyl transfer center. We hypothesized that universally-conserved eIF1A and eIF5B might play a central role in this process. eIF1A resides in the 40S ribosomal A site during initiation and mediates scanning and start codon recognition via its flexible N- and C-terminal tails (*6, 7*). Although the eIF5B-bound 48S initiation complex has never been visualized, a few structures exist for the eIF5B-bound 80S complex (*12-14*). These snapshots suggest that eIF5B core domains (G, II, III) dock onto the 40S subunit outside the A site. Domain IV extends inward to latch onto the acceptor stem of Met-tRNA_i_^Met^, consistent with the proposed role of eukaryote-specific contacts between the C-terminal tail of eIF1A and domain IV of eIF5B in 60S subunit joining (*15-19*). Yet, whether these two factors coordinate to allow ribosomal subunit joining by guiding formation of inter-subunit contacts or by actively remodeling initiation complexes is not known.

To uncover how eIF5B and eIF1A mediate ribosomal subunit joining, we combined real-time single-molecule assays of human translation initiation with targeted structural analyses. We first performed *in vitro* reconstituted experiments where fluorescently-labeled ribosomal subunits, eIF1A, and eIF5B were monitored directly throughout initiation using single-molecule spectroscopy. Our assays defined the timing of association and departure for both proteins, which revealed a transient state where both eIF1A and eIF5B are bound just prior to 60S subunit joining. Using single-particle electron cryo-microscopy (cryo-EM), we characterized structurally this hitherto elusive intermediate, which established the structural basis for eIF5B-catalyzed ribosomal subunit joining.

## RESULTS

### Direct monitoring of human translation initiation

We established a real-time single-molecule assay to monitor human translation initiation directly (**Fig. S1A,B**). β-globin mRNA was tethered to an imaging surface within zero-mode waveguides (ZMWs), which enable four-color fluorescence detection in real time (**Fig. S1C**) (*20*). Saturating concentrations of eIF4 proteins (4A, 4B, 4E, 4G), ATP, and GTP were then added to the tethered mRNA. Separately, the 43S PIC was preassembled and contained 40S subunits labeled with cyanine-3 dye (Cy3, green signal) on the N-terminus of uS19. Upon start of data acquisition at 30 °C, a mixture with 5 nM 43S PIC (by 40S-Cy3), 1 µM eIF5B, and 100 nM 60S subunits labeled with cyanine-5 dye (Cy5, red signal) on the C-terminus of uL18 was added to the imaging surface. When excited by a 532 nm laser, appearance of Cy3 fluorescence indicated that the 40S subunit loaded onto the mRNA. Subsequent appearance of Cy3(donor)-Cy5(acceptor) FRET indicated successful joining of the 60S subunit (FRET efficiency, E_FRET_ = 0.65 ± 0.01) (**Fig. S1D-G**) (*21*). As expected, 40S subunit loading onto mRNA was dependent on the presence of eIF4 proteins (*k*_*on,40S*_ = 0.049 ± 0.002 s^-1^ at 5 nM; ≈10 µM^-1^ s^-1^), whereas 60S subunit joining was dependent on the presence of eIF4 proteins and eIF5B (*k*_*on,60S*_ = 0.033 ± 0.001 s^-1^ at 100 nM) (**Fig. S1H,I**). The mean elapsed time from 40S subunit loading to 60S subunit joining we observed (∼ 30 s) agreed well with initiation timescales obtained in cells (20-30 s) (*22*).

We next tracked fluorescently-labeled eIF5B as it mediates 60S subunit joining. N-terminally truncated eIF5B was labeled on its N-terminus with a Cy3.5 dye via a fused ybbR peptide tag, and the labeled protein was functional in bulk translation (**Fig. S2A,B**). After the eIF4-mRNA complex was tethered in ZMWs, a mixture of the 43S PIC (with 40S-Cy3, green), eIF5B-Cy3.5 (orange), and 60S-Cy5 subunits (red) was added, and fluorescent signals were monitored in real time upon excitation by a 532 nm laser (**Fig. 1A**).

**Figure 1.**
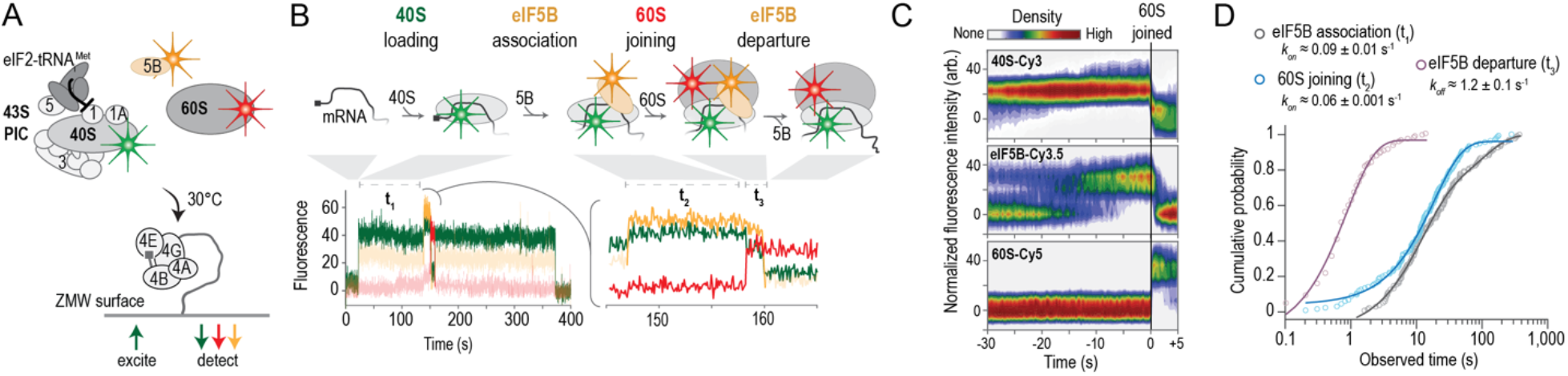
Real-time analysis of eIF5B-mediated ribosomal subunit joining in humans. **A**. Schematic of single-molecule experiment. The 43S PIC (10 nM via 40S-Cy3 subunits, green), 20 nM eIF5B-Cy3.5 (orange), and 100 nM 60S-Cy5 (red) subunit were added to m^7^G-capped β-globin mRNA tethered within ZMWs in the presence of saturating concentrations of eIF4ABGE and 1 mM ATP and GTP at 30 °C. Fluorescence data were acquired at 10 frames per second for 600 s with excitation via a 532 nm laser. **B**. Example single-molecule fluorescence data that depict sequential association of the 40S subunit (green), eIF5B (orange), and the 60S subunit (red). Dwell times correspond to eIF5B association (t_1_), 60S subunit joining (t_2_), and eIF5B departure times from the 80S complex (t_3_). **C**. Density heat map of the normalized 40S-Cy3, eIF5B-Cy3.5, and 60S-Cy5 fluorescence intensities synchronized to joining of the 60S subunit (n = 136). **D**. Observed times for eIF5B association (t_1_), 60S subunit joining (t_2_), and eIF5B departure from the 80S complex (t_3_) at 20 nM eIF5B-Cy3.5 and 100 nM 60S-Cy5 at 30 °C. The 43S PIC was present at 10 nM (via 40S-Cy3). The fast-phase rates are indicated, which corresponds to association (t_1_, t_2_) or dissociation (t_3_) rates (n = 136).

We observed sequential association of the 40S subunit, eIF5B, and the 60S subunit, which was followed by rapid departure of eIF5B (**Figs. 1B,C** and **S2C**). Unlike transient eIF5B sampling previously observed on yeast 48S complexes (*23*), human eIF5B stably associated in a concentration-independent manner with 48S complexes prior to 60S subunit joining (*k*_*on,5B*_ ≈ 0.1 s^-1^ at 20 and 40 nM) (**Figs. 1D** and **S2D**). Further consistent with a slow upstream step that limits its binding, eIF5B associated more rapidly in a concentration-dependent manner (*k*_*on,5B*_ ≈ 20 µM^-1^ s^-1^) with pre-equilibrated 48S complexes (**Fig. S2E-G**), which presumably had progressed past the otherwise rate-limiting step. Conversely, eIF5B association was inhibited ten-fold when departure of eIF2 was prevented using the non-hydrolysable analog GDPNP (**Fig. S2H**). These findings are consistent with a handoff of the Met-tRNA_i_^Met^ acceptor stem from eIF2 to eIF5B, given that both factors bind to the tRNA in this region. The delay from 40S subunit loading until eIF5B association (∼ 10 s) therefore likely represents an upper bound for the integrated timescale of scanning and start codon recognition on β-globin mRNA. Once eIF5B bound, the 60S subunit joined in a concentration- and temperature-dependent manner (*k*_*on,60S*_ ≈ 0.14 ± 0.004 s^-1^ at 200 nM, 30 °C; ≈0.7 µM^-1^s^-1^) (**Figs. 1D** and **S2D,I**). Formation of the 80S initiation complex triggered rapid and temperature-dependent departure of eIF5B (*k*_*off,5B*_ ≈ 1.2 ± 0.1 s^-1^ at 30 °C), which was dependent on GTP hydrolysis (*k*_*off,5B-H706E*_ < 0.01 s^-1^) and the presence of 60S subunits (*k*_*off,5B*_ < 0.01 s^-1^) (**Figs. 1D** and **S2D,I-L**).

We next directly tracked eIF1A-Cy5, labeled at S102 (*24*), throughout translation initiation. We added doubly labeled 43S PICs (40S-Cy3, eIF1A-Cy5), 1 µM unlabeled eIF5B, and 200 nM 60S subunits labeled with Cy5.5 dye (purple) to eIF4-mRNA complexes tethered within ZMWs at 30 °C (**Fig. 2A**). Nearly all recruited 43S PICs began in a 40S(donor)-to-eIF1A(acceptor) FRET state (90 ± 4 %, 158/175; E_FRET_ = 0.42 ± 0.01), as predicted given the proximity of the labeling sites (∼50 Å) (**Figs. 2B** and **S3A-D**). Most complexes were joined by the 60S subunit during the first eIF1A binding event (56 ± 7 %, 88/158) (**Fig. S3B-E**). Complexes with multiple eIF1A binding events also were competent for 60S subunit joining (87/175; k_reass._ ≈ 1.5 ± 0.1 s^-1^ at 45 nM), which suggests the mRNA-loaded complexes can remodel dynamically. While eIF1A bound stably to the 48S complex (k_off_ ≈ 0.075 ± 0.003 s^-1^), the protein departed within a few hundred milliseconds after 60S subunit joining (mean < 0.2 s) (**Figs. 2C,D** and **S3F**). eIF1A departure off the 80S complex was delayed at least 20-fold (mean 3 ± 1.5 s) when GTP hydrolysis by eIF5B was prevented by using GDPNP, consistent with prior findings in yeast (**Fig. 2D**) (*16, 17, 25*). In the absence of eIF1A, the efficiency of 60S subunit joining was reduced 6-fold, and the rare instances of 60S subunit joining occurred at least 4-fold slower (**Fig. S3G,H**). Further confirming that the labeled proteins are functional, the rate of 60S subunit joining (relative to 40S recruitment) was identical in experiments with eIF1A-Cy5 or eIF5B-Cy3.5 (k_on_ ≈ 0.04 ± 0.002 s^-1^ at 200 nM and 30 °C) (**Fig. S3H**), which indicated that the labeled proteins functionally substitute for the unlabeled ones.

**Figure 2.**
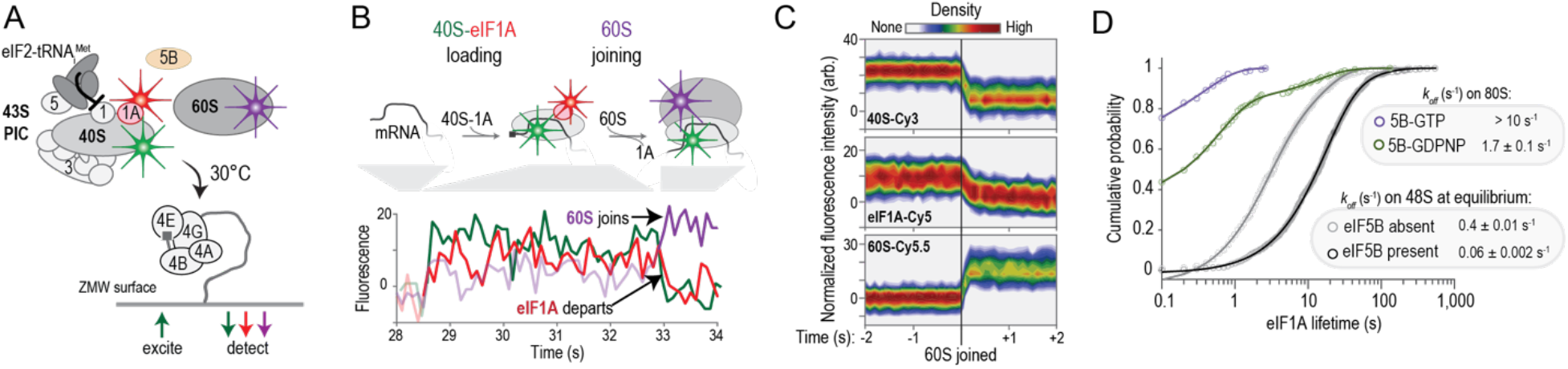
eIF1A resides on initiation complexes until the 60S subunit joins. **A**. Experiment schematic. The doubly labeled 43S PIC (10 nM by 40S-Cy3; eIF1A-Cy5), 1 µM unlabeled eIF5B, and 200 nM 60S-Cy5.5 (purple) subunit were added to β-globin mRNA tethered within ZMWs in the presence of saturating concentrations of eIF4ABGE and 1 mM ATP and GTP at 30 °C. During imaging, eIF1A-Cy5 was present at 4.5-fold molar excess relative to the 40S subunit. Fluorescence data were acquired at 10 frames per second for 600 s with excitation via a 532 nm laser. **B**. Example single-molecule fluorescence data that depict recruitment of a doubly labeled 43S PIC (40S-eIF1A FRET state) followed by 60S subunit joining (40S-60S FRET state) and rapid ejection of eIF1A from the 80S initiation complex. The full fluorescence trace is shown in Fig. S4B. **C**. Density map of the normalized 40S-Cy3, eIF1A-Cy5, and 60S-Cy5.5 fluorescence intensities synchronized to joining of the 60S subunit (n = 175). **D**. Observed eIF1A lifetimes on the 80S initiation complexes in the presence of eIF5B-GTP (purple, n=175) or eIF5B-GDPNP (green, n = 151) or on the pre-equilibrated 48S initiation complex when eIF5B was present (dark grey, n = 589) or absent (light grey, n = 831). Fast-phase dissociation rates are indicated, derived or estimated from fits to double-exponential functions.

eIF1A and eIF5B reside simultaneously on initiation complexes when the 60S subunit joins. Addition of 20 nM eIF5B-Cy3.5 and 10 nM eIF1A-Cy5 to doubly labeled 48S initiation complexes (40S-Cy3 and eIF1A-Cy5) tethered within ZMWs revealed complexes that contained both proteins (**Fig. S4A**). The complexes not only had 40S(Cy3, donor)-to-eIF1A(Cy5, acceptor) FRET signals, but also displayed eIF5B(Cy3.5, donor)-to-eIF1A(Cy5, acceptor) FRET (**Fig. S4B,C**). The latter signal is consistent with the proximity of eIF5B and eIF1A labeling sites predicted by crude structural comparisons (∼70 Å) (**Fig. S4B**). Whereas reassociation rates were similar, eIF1A dissociated from 48S complexes about 7-fold more slowly when eIF5B was present relative to when eIF5B was absent (k_off,fast_ ≈ 0.059 ± 0.002 versus 0.41 ± 0.01 s^-1^) (**Figs. 2D** and **S4D**). Simultaneous addition of doubly labeled 43S PICs (40S-Cy3, eIF1A-Cy5), eIF5B-Cy3.5, and 60S-Cy5.5 subunits to tethered mRNA revealed that nearly all 60S subunit joining events occurred when eIF1A and eIF5B were both present (90 ± 10 %) (**Fig. S4E-H**).

### Structure of an eIF1A-eIF5B-bound initiation complex

To understand how these two proteins collaborate to mediate 60S subunit joining, we used our single-molecule findings to design a vitrification strategy that captured on cryo-EM grids a 48S complex co-bound by eIF1A and eIF5B. Immediately prior to plunge freezing, pre-assembled 48S complexes were incubated with eIF5B for 1 min at 30 °C. At this time point, the 48S complex should be saturated with both eIF1A and eIF5B, and the complex is competent for 60S subunit joining (**Fig. S2D**). Vitrified samples were analyzed by single-particle cryo-EM.

After data collection, image processing, and *in-silico* classifications, we identified a subset of ∼190,000 particles that yielded a high-resolution map after refinement (overall resolution of ∼3.2 Å) (**Fig. S5A,B**). This subset represents ∼84% of all 40S particles, consistent with our prediction above. Robust density for the 40S subunit was visible, as well as density for an aminoacylated tRNA near the ribosomal P site, eIF1A in the A site, and eIF5B (**Fig. 3A**). Peripheral regions of the map are heterogenous, such as the acceptor stem of Met-tRNA_i_^Met^ and eIF5B domains located furthest from the 40S surface, namely domains G and IV. In these flexible areas, local resolution values were in the 4-6 Å range with sufficient quality to place secondary structure elements unambiguously (**Fig. S5C-E**). Density attributable to other eIFs, including eIF3, was absent from cryo-EM maps in this class, even at low map thresholds. The lack of density for eIF3 suggests the factor is highly dynamic on or has departed the initiation complex after eIF2 departure. A small subgroup of 40S particles (∼2100 or 0.65% of the dataset) presented density ascribable to eIF3, but the reduced number of particles precluded further interpretation of this class.

**Figure 3.**
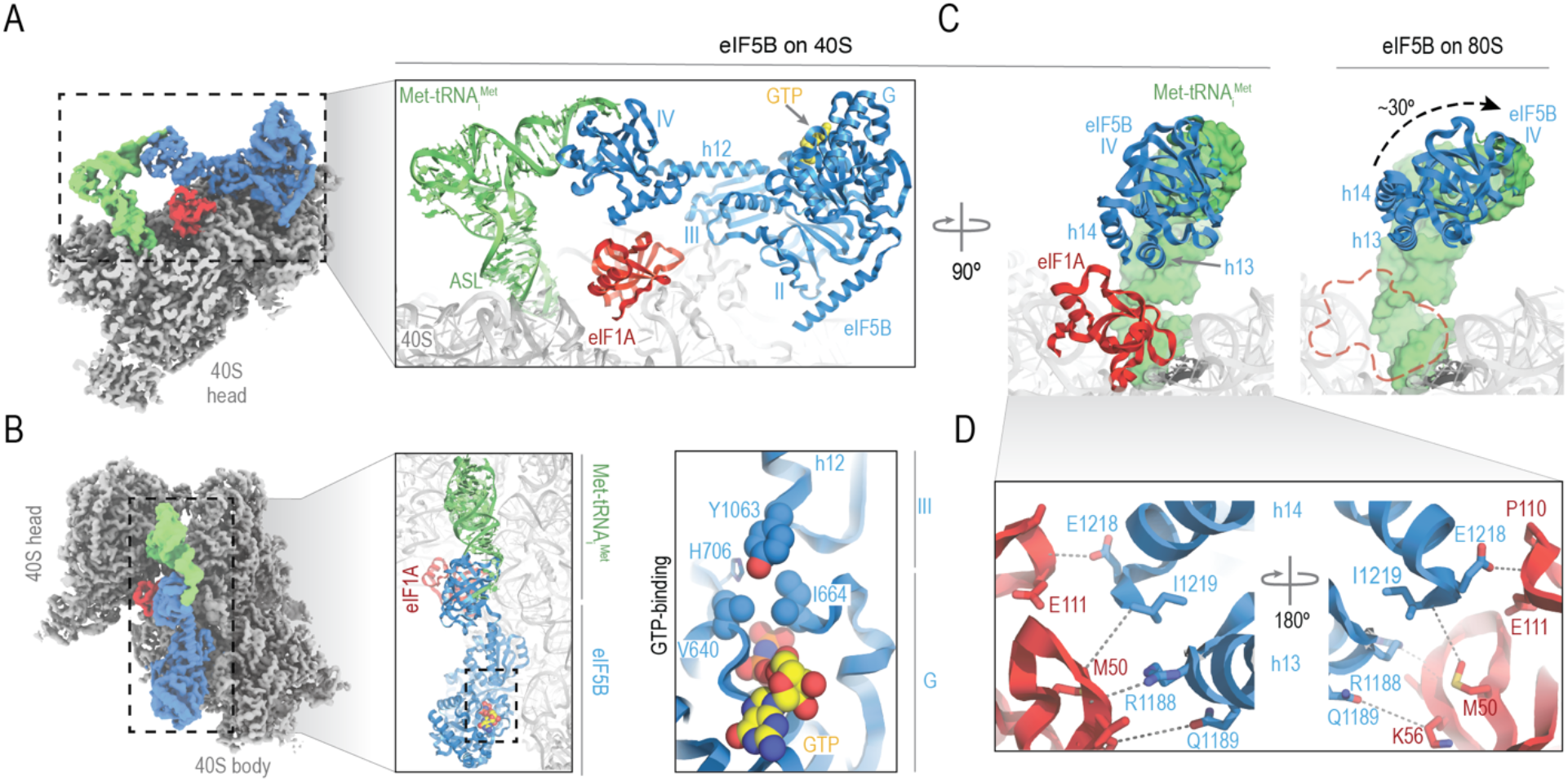
Cryo-EM structure of the eIF5B-bound 48S initiation complex. Post-processed density for the final cryo-EM map (overall resolution 3.2 Å) of the eIF5B-bound 48S-PIC, with 40S subunits colored grey, Met-tRNA_i_^Met^ green, eIF1A red and eIF5B blue. Right, a close-up view of the refined molecular model derived from the cryo-EM data centered around eIF1A, with GTP colored yellow. Domains of eIF5B and regions of Met-tRNA_i_^Met^ are indicated. ASL, anti-codon stem loop; AS, acceptor stem. Left, zenithal view of the cryo-EM map with components colored as in panel A. Middle, view of the molecular model along the acceptor stem axis of Met-tRNA_i_Met. Right, detailed view of the GTP-binding pocket of eIF5B. Tyrosine residue 1063, which belongs to eIF5B domain III, collaborates with G-domain residues V640 and I664, part of the “hydrophobic gate”, in protecting the γ-phosphate of GTP from hydrolysis. **C**. Corresponding views of eIF5B domain IV on the 48S state with eIF1A present (left) and after 60S recruitment with eIF1A ejected (right). Departure of eIF1A allows for a rotational movement of domain IV of eIF5B of approximately 30° towards the 60S, bringing the acceptor stem of Met-tRNA_i_^Met^ close to the peptidyl transfer center (PTC) of the 60S ribosomal subunit. **D**. Zoomed view of the eIF5B-eIF1A interface. Terminal eIF5B α-helices h13 and h14 approach eIF1A, stabilizing weak interactions (>4Å interacting distance) nucleated around eIF1A methionine residue 50 and eIF5B residue isoleucine 1219.

eIF5B density corresponds to the GTP-bound state. The core superdomain of eIF5B, composed of domains G, II, and III, are present in the GTPase binding site on the 40S platform, which represents the largest interaction surface (∼820 Å^2^) in the complex (**Fig. 3A**). The G domain of eIF5B is exposed to solvent, with no direct 40S contacts. It is positioned via interactions with domain III and the base of helix 12. Consistent with a GTP-bound state, switch I of the G domain is ordered and density for the γ-phosphate of GTP is apparent (**Figs. 3B** and **S5E**). The catalytic histidine is maintained away from the γ-phosphate by a steric barrier formed by residues I664 of switch I and V640 of the P loop (**Fig. 3B**, right). These two residues form the prototypical hydrophobic gate that must be pierced by the catalytic histidine to catalyze GTP hydrolysis. The gate is aided by Y1063, as its position at the base of helix 12 interferes with the trajectory the catalytic histidine must follow to reach the γ-phosphate.

Domain IV of eIF5B directly interacts with Met-tRNA_i_^Met^ and eIF1A. Mediated by rigid helix 12 that extends away from the G-II-III superdomain, eIF5B domain IV is latched onto the acceptor stem of Met-tRNA_i_^Met^. Strikingly, relative to its position in 80S initiation complexes (*12-14*), domain IV is rotated about 30 degrees along the helix 12 axis towards the ribosomal A site (**Fig. 3C)**. This rotation positions the two C-terminal alpha-helices of eIF5B (helices 13 and 14) to contact eIF1A directly, which is positioned identical to earlier initiation intermediates (*26-36*) (**Fig. S6A,B**). The interaction interface is relatively small (∼150 Å^2^) and nucleated around a cluster of hydrophobic residues (M50 of eIF1A, and I1219 of eIF5B) (**Fig. 3D**). Whereas the flexible N-terminal tail of eIF1A is disordered, we observed additional density proximal to W1207 on helix 14 of eIF5B that may correspond to I141 of the flexible eIF1A C-terminal tail (**Fig. S6A**). Such contacts are consistent with prior NMR studies on the two proteins in isolation (*37, 38*).

The rotation of eIF5B domain IV induced by eIF1A reorients the Met-tRNA_i_^Met^ configuration to be compatible with 60S subunit joining. We compared our model to two earlier eIF2-bound intermediates, scanning-competent ‘open’ (*34*) or post-scanning ‘closed’ complexes (*33*), and to elongation-competent 80S ribosomes (*39*) (**Fig. S6B**). In our eIF5B-bound complex, the 40S subunit head conformation is similar to both the post-scanning ‘closed’ and elongation-competent conformations (**Fig. S6C**). In contrast, the Met-tRNA_i_^Met^ configuration most closely resembles an elongation state; the anticodon stem loop (ASL) at the 40S P site is nearly identical to the final elongation state, and the acceptor stem and elbow region are almost perpendicular to the ASL (**Fig. 4A**). In this position and mediated by eIF1A, eIF5B domain IV and the acceptor stem are aligned to slot directly into the ∼35 Å channel formed by the central protuberance and helix 69 of the incoming 60S subunit (**Fig. 4B-D**). Thus, our structural analyses suggest that the 40S–eIF1A– eIF5B–acceptor-stem network of interactions are needed to funnel multiple conformations of Met-tRNA_i_^Met^ towards a position compatible with 60S subunit joining.

**Figure 4.**
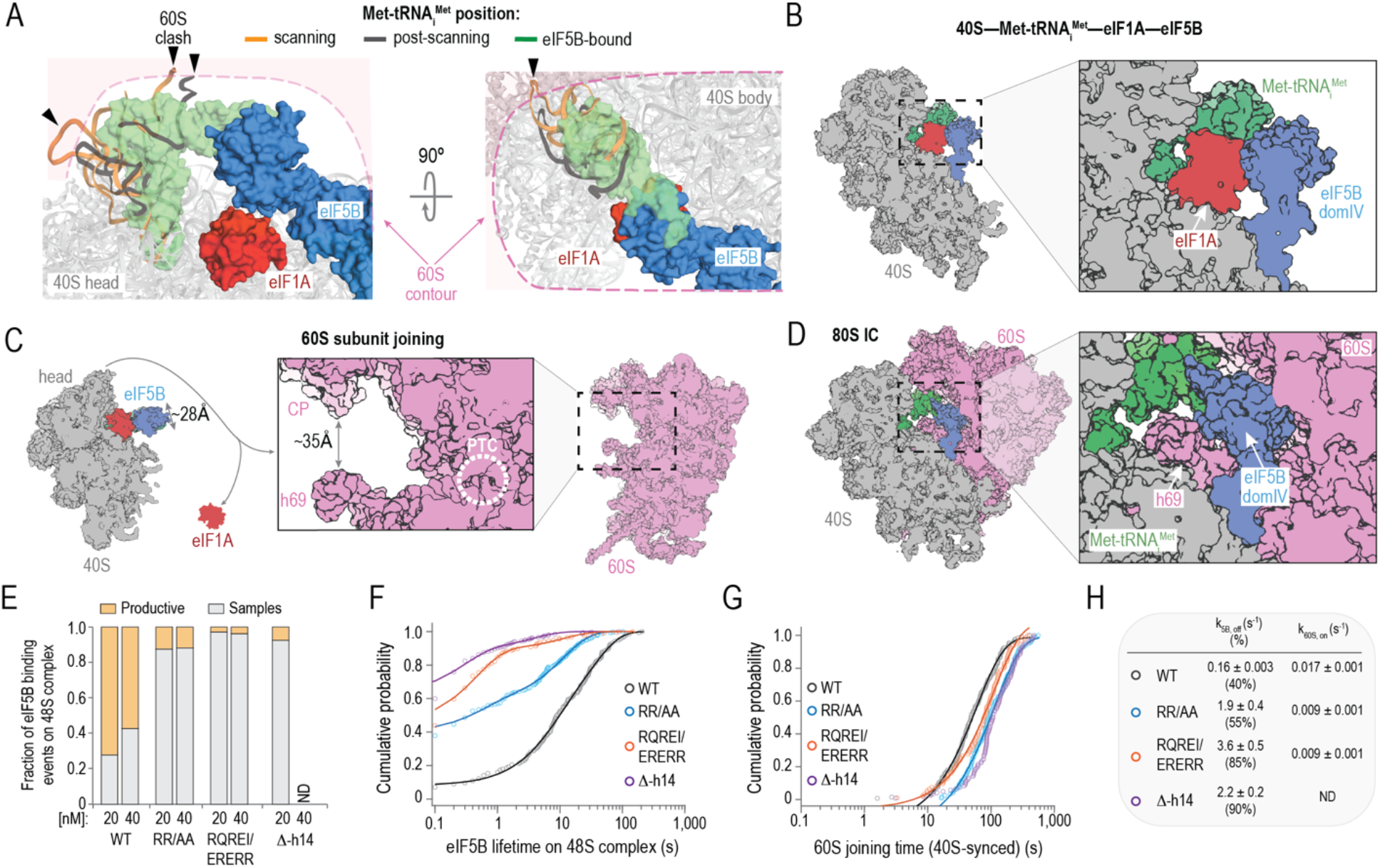
eIF5B contacts with eIF1A and Met-tRNA_i_^Met^ mediate 60S subunit joining. **A**. Comparison of Met-tRNA_i_^Met^ conformations in scanning-competent (orange), post-scanning (grey), and eIF5B-bound 48S (green) complexes. eIF5B reorients the elbow and acceptor stem region of Met-tRNA_i_^Met^ to avoid steric clashes (black arrowheads) with the incoming 60S subunit. **B-D**. Structural models for how eIF1A and eIF5B domain IV orient the acceptor stem of Met-tRNA_i_^Met^ to slot into the channel formed by the central protuberance (CP) and helix 69 of the incoming 60S subunit. As it joins, helix 69 ejects eIF1A from the complex. **E**. Fraction of observed eIF5B binding events on the 48S complex that were samples or productive, defined as whether the event was distinct from or overlapped with 60S subunit joining, respectively, for wild-type and the indicated mutant eIF5B proteins. **F**. Observed lifetimes of the indicated eIF5B proteins on the 48S complex at 20 nM at 30 °C. n = 156, 251, 196, and 320. **G**. Observed 60S joining times, quantified relative to 40S loading, when the indicated eIF5B protein was present at 20 nM at 30 °C. n = 133, 123, 93, and 121. **H**. Summary of eIF5B dissociation rates from and 60S subunit association rates with the 48S initiation complex. Only fast phases rates are shown here, see Fig S7 for all rates.

We used our single-molecule assays to assess the role of residues in eIF5B that our cryo-EM-derived model suggest mediate contacts with Met-tRNA_i_^Met^ or eIF1A in translation initiation. Contacts with the Met-tRNA_i_^Met^ acceptor stem were disrupted by substitution of conserved R1105 and R1174 with alanine (RR/AA) (**Fig. S7A-C**). Contacts with eIF1A were disrupted either by truncation of eIF5B prior to helix 14 (Δ-h14) or five targeted substitutions of residues in helices 13 and 14 that invert surface contacts with eIF1A (RQREI/ERERR) (**Fig. S7A-C**). After the eIF4-mRNA complex was tethered in ZMWs, a mixture of the 43S PIC (with 40S-Cy3, green), wild-type or mutant eIF5B-Cy3.5 (orange), and 60S-Cy5 subunit (red) was added and fluorescent signals were monitored in real time (**Fig. S7D-F**). Relative to the wild-type protein, the three mutant eIF5B proteins were much more likely (5-7 fold) to rapidly sample the 48S complex (nearly 100-fold decreased lifetimes) (**Fig. 4E,F**). The destabilized mutant eIF5B proteins reduced the efficiency of 60S subunit joining, and when the 60S did join, it occurred at least 2-3 fold more slowly than with the wild-type protein (**Fig. 4G,H** and **S7H-K**). Thus, the eIF5B–eIF1A–40S–Met-tRNA_i_^Met^ interactions stabilize and remodel the 48S initiation complex to mediate 60S subunit joining.

## DISCUSSION

Our single-molecule and structural results suggest a model for how eIF1A and eIF5B mediate human ribosomal subunit joining (**Fig. 5** and **Movie 1**). The eIF1A–40S subunit complex – presumably as the 43S PIC – loads onto the mRNA. eIF1A remains stably bound in the A site throughout scanning and start codon recognition, consistent with its key roles in those steps. Our structure indicates that the flexible N-terminal tail of eIF1A is absent from the P site after eIF2 departs the complex, which we estimate occurs in about ∼10 s on β-globin mRNA. eIF2 departure also enables eIF5B to stably bind the complex and form the 40S late initiation complex (40S LIC). Stable eIF5B association with the 40S LIC very likely provides the molecular basis for how eIF5B prevents reassociation of eIF2-GDP with Met-tRNA_i_^Met^ (*40*). Furthermore, relative to yeast (*23*), the enhanced stability of human eIF5B on the 40S LIC provides new insight to understand how increased eIF5B abundance drives aberrant translation in lung adenocarcinomas (*41*).

**Figure 5.**
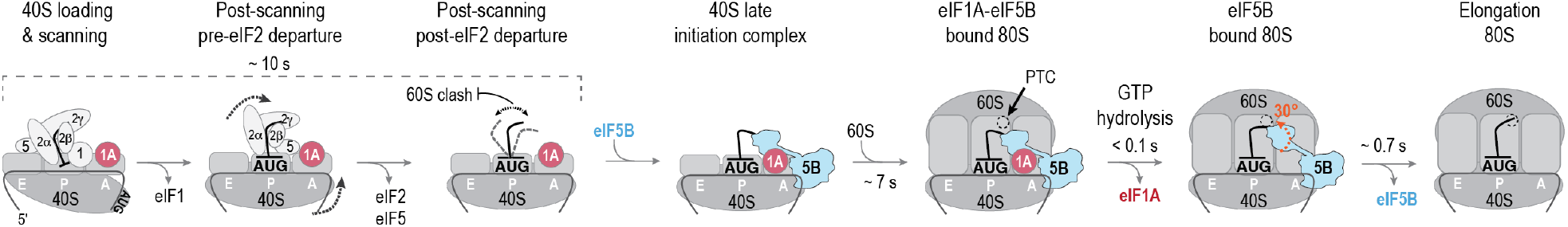
Proposed model for how human eIF1A and eIF5B collaborate to orient Met-tRNA_i_^Met^ properly and catalyze ribosomal subunit joining, as illuminated by our single-molecule dynamics and structural results.

For eIF5B to bind stably to the 40S LIC, domain IV of the protein must contact both eIF1A and Met-tRNA_i_^Met^. Loss of contacts with either led to a 100-fold decrease in eIF5B lifetime on the complex and inhibited 60S subunit joining. Thus, multiple relatively weak interactions likely stabilize this conformationally-flexible GTPase on the initiation complex. Intriguingly, previous structural analyses of the protein in isolation or bound to eIF1A have suggested that eIF5B adopts both extended and compact conformations, with domain IV either extended away from or adjacent to the G-II-III superdomain (*37, 38, 42-44*). In the context of initiation, contacts with eIF1A may enhance or stabilize extension of domain IV away from the G-II-III superdomain of eIF5B, as the compact conformation would clash sterically with eIF1A in the ribosomal A site.

Eukaryote-specific contacts between eIF1A and eIF5B reconfigure the Met-tRNA_i_^Met^ acceptor stem, driven by rotation of eIF5B domain IV along the helix 12 axis. This surprising rotation – rather than a rigid, swing-like movement predicted by studies in prokaryotes (*45*) and archaea (*42*) – funnels the complex into a conformation that slots directly into the channel formed by the central protuberance and helix 69 of the 60S subunit. Formation of the 80S ribosome triggers hydrolysis of GTP by eIF5B to accelerate ejection of eIF1A by helix 69 of the 60S subunit (within ∼100 ms), likely by accelerating back-rotation of the 40S subunit from a semi-rotated state to a classical non-rotated state, similar to prokaryotes (*46-48*). Indeed, inter-subunit back-rotation moves the apical loop of helix 69 by ∼16 Å further into the ribosomal A site, where eIF1A resides. eIF1A departure frees eIF5B domain IV and the acceptor stem to rotate ∼30 degrees toward the peptidyl transfer center. The rotation poises the aminoacylated end of the tRNA for its final accommodation upon eIF5B departure, which occurrs within another few hundred milliseconds and is dependent on GTP hydrolysis. Dynamic rotation of eIF5B domain IV mediated by eIF1A therefore plays a key role in the late steps of translation initiation.

Collectively, our findings suggest that the eIF1A–eIF5B–Met-tRNA_i_^Met^ network of interactions serves as a final fidelity checkpoint to ensure that initiation complexes are competent for the 60S subunit to join. They also demonstrate that eIF5B, mediated by contacts with eIF1A, actively remodels the 40S LIC to facilitate 60S subunit joining, rather than serving simply as a chaperone of inter-ribosomal subunit contacts. Almost immediately after formation of the 80S initiation complex, GTP hydrolysis by eIF5B catalyzes a series of events that clears the ribosomal A site and positions the aminoacylated end of Met-tRNA_i_^Met^ in the peptidyl transfer center of the ribosome, which commits the ribosome to synthesize a protein.

## Acknowledgements

We are grateful to Michael Lawson, Jan Carette, and other members of the Puglisi and Carette labs for helpful guidance, discussions, and feedback. We thank Peter Sarnow and the Sarnow lab for sharing cell culture equipment. Some of this work was performed at the Stanford-SLAC Cryo-EM Facilities, supported by Stanford University, SLAC and the National Institutes of Health S10 Instrumentation Programs. The content is solely the responsibility of the authors and does not necessarily represent the official views of the National Institutes of Health.

## Author contributions

Conceptualization: CPL, ISF, JDP

Methodology: CPL, RG, MS, CA, JW, BSS, EM, TED, CSF, ISF, JDP

Resources: CPL, RG, MS, CA, NV, CSF

Investigation: CPL, RG, EM, ISF

Visualization: CPL, ISF, JDP

Funding acquisition: TED, CSF, JDP

Project administration: TED, CSF, ISF, JDP Supervision: TED, CSF, ISF, JDP

Writing – original draft: CPL, ISF, JDP

Writing – review & editing: CPL, RG, MS, CA, JW, TED, CSF, ISF, JDP

## Data and materials availability

All single-molecule data are included in the manuscript.

## MATERIALS AND METHODS

### eIF5B cloning

N-terminally truncated (587-1220) with an N-terminal ybbR-tag eIF5B and relevant mutants were purchased as geneblocks from IDT and cloned into a vector purchased from the UC Berkeley QB3 MacroLab (vector 1B) using their standard protocol. Contacts with Met-tRNA_i_^Met^ were disrupted by R1105A and R1174A substitution. Contacts with eIF1A were disrupted by truncation of eIF5B at Q1203 (Δ-helix 14) or five substitutions that inverted properties along the interaction interface (R1188E, Q1189R, R1199E, E1218R, I1219R). All sequences were verified by Sanger sequencing. The resulting plasmids encoded eIF5B(587-1220) tagged on the N-terminus with a 6-histidine tag fused to maltose-binding protein (MBP) followed by a TEV protease cleavage site, a short NA linker, and the ybbR peptide tag (NH_2_-6His–MBP–TEV–NA–ybbR– eIF5B-COOH).

### β-globin mRNA

A synthetic DNA was purchased from IDT that contained a 5’ flanking sequence (pUC19 backbone sequence), a T7 promoter (TAATACGACTCACTATAG), the human β-globin transcript 1 (NM_000518.5, 628 nts), eight (8) non-templated adenosines, and a 3’ flanking sequence (pUC19 backbone sequence).

Linear DNA templates were amplified using NEB Phusion polymerase and a 5’ primer (TGACCATGATTACGCCAAGC) that annealed upstream of the T7 promoter and a 3’ primer (TTTTTTTTTTTTTTTTTTTTTTTTTTTTTTttgcaatgaaaataaatgttttttattagg) that annealed to the 3’ end of β-globin transcript and encoded a poly(A)_30_ tail. The mRNA was in vitro transcribed using the T7 Megascript kit (Invitrogen, AM1334) and standard conditions. Transcribed RNAs were purified using the GeneJET RNA Purification Kit (ThermoFisher, #K0732). The 3’-terminus was biotinylated by potassium periodate oxidation followed by reaction with biotin-hydrazide, as described (*49*). The 5’-terminus was capped with a type I 7-methylguanosine cap (m^7^G) using the Vaccinia Capping System (NEB, M2080S) and 2’-O-methyltransferase (NEB, M0366), following the one step protocol. 5’-capped and 3’-biotinylated β-globin mRNA was purified using the GeneJET RNA Purification Kit and stored at -20 °C until use.

### Purification and fluorescent labeling of ybbR-tagged eIF5B

All wild-type and mutant ybbR-eIF5B expression plasmids were transformed into Rosetta2 cells purchased from the UC Berkeley QB3 MacroLab and grown overnight at 37 °C on LB agar plates supplemented with 50 µg/mL kanamycin. Liquid cultures of single colonies were grown to OD_600_ ≈ 0.5 at 37 °C in LB supplemented with kanamycin. Cultures were shifted to 18 °C for 30 minutes, 0.5 mM IPTG was added, and cultures were grown for 16-20 h at 18 °C. Cells were harvested by centrifugation at 5,000 x *g* for 15 min at 4 °C in a Fiberlite F9 rotor (ThermoFisher, cat. # 13456093). Cells were lysed by sonication in lysis buffer (20 mM Tris-HCl pH 8.0, 300 mM NaCl, 10% (v/v) glycerol, 20 mM imidazole, and 5 mM β-mercaptoethanol), and lysates were cleared by centrifugation at 38,000 x *g* for 30 min at 4 °C in a Fiberlite F21 rotor followed by filtration through a 0.22 µm syringe filter. Clarified lysate was loaded onto a Ni-NTA gravity flow column equilibrated in lysis buffer, washed with 20 column volumes (CV) of lysis buffer, 20 CV of wash buffer (20 mM Tris-HCl pH 8.0, 1 M NaCl, 10% (v/v) glycerol, 40 mM imidazole, and 5 mM β-mercaptoethanol), and 10 CV of lysis buffer. Recombinant proteins were eluted with five sequential CV of elution buffer (20 mM Tris-HCl pH 8.0, 300 mM NaCl, 10% (v/v) glycerol, 300 mM imidazole, and 5 mM β-mercaptoethanol). Fractions with recombinant protein were identified by SDS-PAGE analysis. The relevant fractions were dialyzed overnight at 4 °C into TEV cleavage buffer (50 mM HEPES-KOH pH 7.5, 200 mM NaCl, 10% (v/v) glycerol, and 1 mM DTT) in the presence of excess TEV protease. Fluorescent labeling via the ybbR tag was performed essentially as described (*49, 50*). Briefly, ∼10 µM ybbR-eIF5B was supplemented with 10 mM MgCl_2_ and incubated at 37 °C for 90 min in the presence of 2-4 µM Sfp synthase enzyme and 20 µM of Cy3.5-CoA substrate. Free dye was removed via purification over 10DG-desalting columns (Bio-Rad, cat.# 7322010) equilibrated in TEV cleavage buffer supplemented with 20 mM imidazole. TEV protease, Sfp synthase, and the cleaved 6His tag were removed via a subtractive Ni-NTA gravity column equilibrated in TEV buffer, with the flow-through collected. Labeled proteins were purified using size exclusion chromatography (SEC) on a Superdex 75 or 200 column (23 mL) equilibrated in SEC (storage) buffer (20 mM HEPES-KOH pH 7.5, 150 mM KOAc, 10% (v/v) glycerol, and 1 mM DTT).

Fractions containing eIF5B proteins were concentrated using a 30 kD MWCO Amicon Ultra centrifugal filter, aliquoted, flash frozen on liquid N_2_, and stored at -80 °C.

### *in vitro* extract-based translation assays

The nanoluciferase GAPDH reporter mRNA was prepared as described (*21*). Standard HeLa cell-free translation (ThermoFisher, #88882) reactions were programed with a final mRNA concentration of 100 nM and 840 nM of the indicated eIF5B protein. An equal volume of eIF5B SEC (storage) buffer (without protein) was added to a control reaction to account for buffer effects on IVT activity. IVT reactions were incubated at 37°C for 45 min and then immediately transferred to an ice water bath and diluted 1:1 with cold Glo Lysis Buffer (Promega, #E2661). All samples were brought to room temperature and mixed with a 1:1 volume of nGlow solution (Promega, #N1110). Samples (90% of total volume) were loaded into non-adjacent wells of a 384-well plate. Sample luminescence was measured 11 min post nGlow solution addition using a BioTek Neo2 multi-mode plate reader (25°C, 114LUM1537 filter, gain of 135). Luminescence signal was monitored for an additional 30 min at 3 min intervals to verify luminesce signal of all samples decayed at the same rate. Prism8 (Graphpad) was used for all analyses.

### Purification of human eIFs, Met-tRNA_i_^Met^, and ribosomal subunits

Recombinant eIF1 (*51*), eIF1A (*51*), eIF3j (*51*), eIF4AI (*51, 52*), eIF4E (*52, 53*), and eIF5 (*54*) proteins were purified as described. Recombinant eIF4B was purified essentially as described (*51, 52*), except cells were lysed solely by sonication and a subtractive Ni-NTA step was added prior to ion exchange chromatography (heparin). Coding sequence for eIF4G(165-1599) was inserted into 6His-tagged pFASTBAC1 vector and purified similar to a described protocol (*53*), with the following modifications: 15 µM leupeptin was added to growth media 24 hrs prior to harvest, cells were lysed by sonication, and a subtractive Ni-NTA step was added, which was followed by ion exchange chromatography (heparin) and concentration into SEC buffer with 300 mM KOAc. Native human eIF2 and eIF3 (lacking 3j) were purified as described (*51, 55*). Human Met-tRNA_i_^Met^ was in vitro transcribed and aminoacylated as described (*21*). eIF1A(S102C) was purified and fluorescently labeled with Cy5 dye as described (*24*). Human 40S and 60S ribosomal subunits were purified from the appropriately edited HEK293T cell lines and labeled with fluorescent dyes as described (*21, 49*).

### ZMW-based single-molecule spectroscopy

All real-time imaging was conducted using a modified Pacific Biosciences RSII DNA sequencer, which was described previously (*56*). All experiments were performed at 30 °C (unless otherwise noted) using a 532 nm excitation laser at 0.32 µW/µm^2^, which directly excited Cy3 and Cy3.5 dyes. Cy5 and Cy5.5 dyes were excited via FRET as indicated. Four-color fluorescence emission (Cy3, Cy3.5, Cy5, and Cy5.5) was detected at 10 frames per second for 600 s. ZMW chips were purchased from Pacific Biosciences. Prior to imaging, all ZMW chips were washed with 0.2% Tween-20 and TP50 buffer (50 mM Tris-OAc pH 7.5, 100 mM KCl). Washed chips were coated with neutravidin by a 5 min incubation with 1 µM neutravidin diluted in TP50 buffer supplemented with 0.7 mg mL^-1^ UltraPure BSA and 1.3 µM of pre-annealed DNA blocking oligos (CGTTTACACGTGGGGTCCCAAGCACGCGGCTACTAGATCACGGCTCAGCT) and (AGCTGAGCCGTGATCTAGTAGCCGCGTGCTTGGGACCCCACGTGTAAACG). The imaging surface then was washed with TP50 buffer at least four times.

#### Real time initiation single-molecule assays

For all single-molecule assays, the ‘initiation reaction buffer’ was: 20 mM HEPES-KOH, pH 7.3, 70 mM KOAc, 2.5 mM Mg(OAc)_2_, 0.25 mM spermidine, 0.2 mg mL^-1^ creatine phosphokinase, 1 mM ATP•Mg(OAc)_2_, and 1 mM GTP•Mg(OAc)_2_. The ‘imaging buffer’ was the initiation reaction buffer supplemented with casein (62.5 µg mL^-1^) and an oxygen scavenging system (*57*): 2 mM TSY, 2 mM protocatechuic acid (PCA), and 0.06 U/µL protocatechuate-3,4-dioxygenase (PCD).

To prepare the eIF2–GTP–Met-tRNA_i_^Met^ ternary complex (TC), 3.3 µM eIF2 was incubated in initiation reaction buffer (excluding ATP•Mg(OAc)_2_) for 10 min at 37 °C to saturate eIF2 with GTP. The eIF2–GTP complex then was incubated with 2.3 µM Met-tRNA_i_^Met^ for 5 min at 37 °C to form TC. In a few instances, GTP was replaced with a non-hydrolyzable analog (GDPNP) during formation of TC, which prevents GTP hydrolysis and subsequent departure of eIF2 from the initiation complex.

To prepare the 43S pre-initiation complex (43S PIC), 1 µM eIF1, 1 µM eIF1A, 500 nM TC (by eIF2), 1.2 µM eIF5, 400 nM eIF3, 1.2 µM eIF3j, and 240 nM 40S-Cy3 subunits were incubated for 5 min at 37 °C in initiation reaction buffer. In experiments with doubly labeled 43S PICs (40S-Cy3, eIF1A-Cy5), unlabeled eIF1A was substituted for an equal concentration (by Cy5) of eIF1A(S102C)-Cy5.

In all full real-time single-molecule initiation assays, β-globin mRNA with a 5’ m^7^G cap, poly(A)_30_ tail, and 3’-terminal biotin moiety was tethered to a prepared and neutravidin-coated ZMW imaging surface.

Immediately after tethering and washing the surface with imaging buffer, a mixture of 2 µM eIF4A, 440 nM eIF4B, 260 nM eIF4G, and 320 nM eIF4E in 20 µL of imaging buffer was added to the surface. The surface and reaction mixture were incubated at room temperature for 5-10 minutes as the instrument initialized. At the start of data acquisition, a 20 µL mixture of 10-20 nM 43S PIC (by 40S-Cy3), 2 µM eIF5B, and 200-400 nM 60S-Cy5/5.5 subunits in imaging buffer was added to the surface. In experiments with fluorescently-labeled eIF5B, unlabeled eIF5B was replaced with 40-80 nM eIF5B-Cy3.5. Importantly, addition of this second reaction mixture to the surface doubled the total reaction volume (to 40 µL, since 20 µL with eIF4 proteins was already present); thus, final concentrations of all components during data collection are halved from the values reported above. Exact final concentrations for labeled components in particular figures are reported in the legends and text.

In a few instances, the 48S initiation complex was pre-equilibrated and tethered to the imaging surface. To do this, 90 nM 43S PIC (by 40S-Cy3), 90 nM of β-globin mRNA, 2 µM eIF4A, 1.1 µM eIF4B, 650 nM eIF4G, and 800 nM eIF4E was incubated for 15 min at 37 °C in initiation reaction buffer. The resulting complex was diluted 1:12.5, immediately added to a prepared and neutravidin-coated ZMW imaging surface, and incubated for 10 min at room temperature to tether the complex to the surface. After washing the surface with imaging buffer, a 20 µL mixture of 1 µM eIF1A, 2 µM eIF4A, 440 nM eIF4B, 260 nM eIF4G, and 320 nM eIF4E in 20 µL of imaging buffer was added to the surface. At the start of data acquisition, a 20 µL mixture of 40-80 nM eIF5B ± 200 nM 60S-Cy5 subunits in imaging buffer was added to the surface. In experiments with labeled eIF1A, a 20 µL mixture with 20 nM eIF1A-Cy5 and 40 nM eIF5B-Cy3.5 was added to the imaging surface with a tethered and doubly labeled 48S PIC (where unlabeled eIF1A was replaced with labeled eIF1A). As above, final concentration of all components during data collection are halved from the reported values.

To probe the role of GTP hydrolysis by eIF5B, the GTP in the imaging buffer was replaced with an equal concentration of the non-hydrolyzable analog GDPNP. In this experimental setup, TC would remain in the GTP-bound state, as eIF2 required a specific exchange factor to exchange GTP/GDP.

#### Data analysis

Experimental movies that captured fluorescence intensities over time were processed using MATLAB as described previously (*20, 21, 49*). In all analyses, ZMWs with the desired fluorescence signals were identified by filtering for the desired signals.

To determine *E*_*FRET*_, fluorescence data from individual ZMWs with the desired fluorescence signals (e.g., 40S-60S FRET) were background corrected using SPARTAN (*58*), with single molecules indicated by single-step photobleach events of the donor fluorophore. FRET on and off states were assigned automatically using vbFRET (*59*), which were visually inspected and manually corrected as needed. An *E*_*FRET*_ threshold of 0.1 was used in all experiments. *E*_*FRET*_ was defined as:

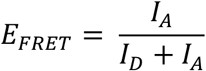

Where *I*_*D*_ and *I*_*A*_ represent fluorescence intensities of the donor and acceptor fluorophores. *E*_*FRET*_ values observed across all events and molecule were binned (50 bins, -0.2 to 1.2) and fit to gaussian functions to determine the mean and standard deviations.

To determine association and dissociation kinetics, binding events of individual components (e.g., 40S, eIF5B, and 60S) were assigned manually based on the appearance and disappearance of the respective fluorescence signals. The observed times for an event to occur from 100-200 individual molecules (i.e., ZMWs) were used to calculated cumulative probability functions of the observed data (cdfcalc, MATLAB), which were fit to single- or double-exponential functions in MATLAB (cftool, non-linear least squares method) as described (*20, 60*). All derived association rates, median association times, lifetimes, and the number of molecules examined are reported in the relevant figures and legends. The exponential function was defined as:

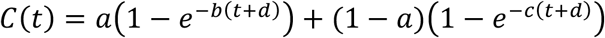

where *t* is time (in s), *b* and *c* rates, and *d* an adjustment factor. If a parameter yielded a phase that represented less than 10% of the population, only a single phase was used to derive the respective rate and reported (i.e., single-exponential function).

#### Statistical analyses

To calculate errors for the efficiency of a given binding event, bootstrap analyses (n = 10,000) were performed to calculate 95% confidence intervals (C.I.) for the observed proportions using R (mosaic library, rflip approach). To calculate errors for median association times and lifetimes, bootstrap analyses (n = 10,000) were performed to calculate 95% C.I. of the observed median using MATLAB (bootstrp and bootci functions). Reported errors for derived rates represent 95% C.I. yielded from fits to linear, single-exponential, or double-exponential functions, as indicated.

### Formation of the eIF5B-bound 48S initiation complex for cryo-EM analysis

The eIF2–GTP–Met-tRNA_i_^Met^ ternary complex was formed as above. To prepare the 43S PIC, 1.5 µM eIF1, 1.5 µM eIF1A, 1 µM TC (by eIF2), 1.5 µM eIF5, 800 nM eIF3, and 600 nM 40S-Cy3 subunits were incubated for 5 min at 37 °C in initiation reaction buffer. The 43S PIC was incubated on ice. In parallel, 200-mesh Quantifoil R2/1 grids (Electron Microscopy Sciences, Q250AR1) were glow-discharged for 25 s in a PELCO EasiGlow glow discharger (Ted Pella, Inc., conditions: negative charge, 15 mA, 0.4 mBar, 25 s). After grids were prepared, 300 nM 43S PIC (by 40S), 500 nM of β-globin mRNA, 1 µM eIF4A, 1 µM eIF4B, 650 nM eIF4G, and 800 nM eIF4E were incubated for 3 min at 37 °C in initiation reaction buffer. Immediately after, 1 µM eIF5B was added and the resulting reaction mixture was incubated for ∼55 s at 30 °C. A 3 µL sample from the mixture was applied to each grid at 21 °C and 95% humidity. The total time elapsed after eIF5B addition until application to the grid was ∼60 s, and the final concentration of 40S subunit was 250 nM. The sample was vitrified by plunging into liquid ethane after 2.5 s blotting using a Leica EM GP (Leica Microsystems) plunger.

### Cryo-EM data collection, image processing, and structure determination

Vitrified grids were screened for ice thickness and intactness on a Glacios microscope (ThermoFisher) equipped with a Gatan K2 Summit direct electron detector. High-resolution data were collected on a Titan Krios equipped with a BioQuantum-K3 imaging system operated in super-resolution mode with pixel size 0.42Å/pixel and fluence of 20 e/A^2^/s. The defocus range was set between -0.5 and -2 micrometers. Image processing was done using Relion3.1 (*61*) with ctf estimation via a wrapper to CtfFind4 (*62*). After 2 times binning in motion correction and ctf estimation, a laplacian autopicking job identified 2 million particles in the dataset. Initial particle extraction was performed with a binned size of 8 and a 2D classification job (200 classes, tau fudge 4 and the subset option deactivated) was used to discard bad particles or poorly aligned ones. The selected particles were subjected to a 3D refinement job using as initial model a volume obtained by stochastic gradient descent (SGD). After 3D refinement, particles were re-extracted at bin 8 with the recenter option activated. Next, a 3D classification job using a soft mask around the factors and the tRNA and without alignment (20 classes, tau-fudge parameter set to 3) found a major population with clear features for the tRNA and the factors, particles belonging to this class (∼109,000) were selected, re-extracted to bin 2 and re-refined at this pixel size. This reconstruction reached ∼4 Å resolution which was further improved by refinement of the ctf parameters and Bayesian polishing of the beam induced particle movements. The final subset of “shiny” particles was re-refined and reached an overall resolution of 3.2 Å after post-processing.

Map exploration revealed features compatible with the reported resolution, like clear side chain density, especially for tyrosine, phenylalanine, leucine and tryptophan residues. Final postprocessed maps were used for model building in coot (*63*). Good general density, especially for bulky residues, allowed proper tracing of the polypeptide of human eIF5B and eIF1A. Modelling of eIF5B was aided by the AlphaFold prediction (*64*), which domain wise, fit the experimental maps accurately. Only minor manual, rigid-body adjustments of individual domains were necessary to fully account for the experimental density prior to stereochemical model refinement. The final model was refined in real space with phenix imposing secondary structure and Ramachandran restrains (*65*). This refined model was checked in coot, corrected in problematic areas, and re-refined in the reciprocal space with Refmac, using secondary structure restrains computed in ProSmart (*66*). The quality of the final model was monitored by Molprobity (*67*). Pymol, Chimera and ChimeraX (*68*) were used for analysis of the structure and figure making.

**Supplementary Figure 1.**
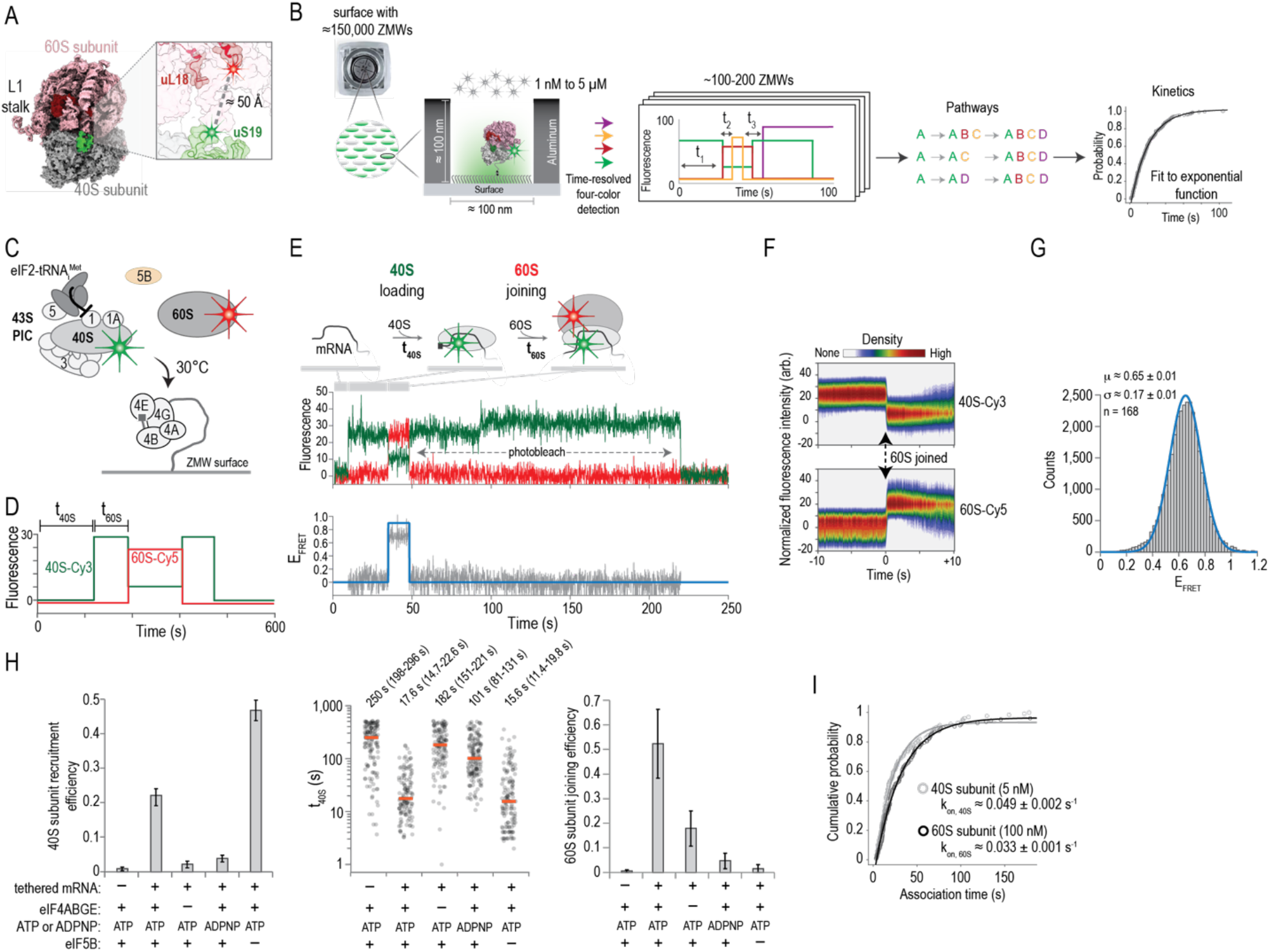
Real-time monitoring of human ribosomal subunit recruitment to β-globin mRNA. **A**. Human 40S and 60S ribosomal subunits were labeled with fluorescent dyes on the N-terminus of uS19 or C-terminus of uL18, respectively. These labeling positions yield an inter-subunit FRET signal upon formation of translation-competent 80S ribosomes. The structural model was obtained from PDB: 4UG0. **B**. A real-time single molecule fluorescence assay using zero-mode waveguides (ZMWs) on a custom PacBio RSII DNA sequencer. Components of interest are tethered to the imaging surface within individual ZMWs, and reaction components are added directly to the surface. Time-resolved, real-time 4-color fluorescence emission was monitored across ∼150,000 ZMWs after excitation with a 532 nm laser. The order and time elapsed between relevant fluorescence signals (e.g., green & orange) were determined in ∼100-200 ZMWs with the desired signals. Rates of association and dissociation were determined using probability based statistical models; Cumulative distribution functions of the observed times were calculated and subsequently fit to exponential functions to yield association dissociation rates, as appropriate. **C**. Schematic of a real-time single-molecule assay to monitor ribosomal subunit joining on tethered β-globin mRNA (‘tethered mRNA’). β-globin mRNA with a 5’ m^7^G cap, poly(A)_30_ tail, and 3’-terminal biotin moiety was tethered to the neutravidin-coated ZMW imaging surface. Immediately after tethering, eIFs 4A, 4B, 4G, and 4E were added. After start of data acquisition and excitation via the 532 nm laser at 30 °C, the 43S PIC (5 nM via the labeled 40S-Cy3), unlabeled eIF5B (1 µM), 60S-Cy5 subunits (100 nM), and 1 mM of ATP and GTP were added. Unlabeled 43S PIC components were present at ≥ 1.5-fold excess during each step of the experiment. Fluorescence data were acquired for 600 s. **D**. Cartoon schematic of theoretical single-molecule fluorescence data where 40S and 60S subunits were recruited to an mRNA to form 80S ribosomes. The 40S subunit association time (t_40S_) was defined as the time elapsed from addition of the 43S PIC until appearance of the Cy3 signal. The 60S subunit association time (t_60S_) was defined as the time elapsed from 40S subunit association until appearance of the 40S(Cy3,donor)-60S(Cy5,acceptor) FRET signal. **E**. Example single-molecule fluorescence trace with FRET efficiency (E_FRET_) plot. Recruitment of the 40S subunit (as the 43S PIC) was indicated by a burst of Cy3 (green) fluorescence intensity. 60S subunit joining was indicated by appearance of the 40S-60S FRET signal. **F**. Density maps of normalized 40S-Cy3 and 60S-Cy5 fluorescence intensities post-synced to 60S subunit joining (n = 128). As in the individual trace shown in panel D, joining of the 60S subunit led to an anti-correlated decrease in Cy3 and increase in Cy5 signals, indicative of intersubunit FRET when the 80S ribosome formed. **G**. Histogram and single gaussian function fit (line) of the observed E_FRET_ in the real-time initiation assay. The mean E_FRET_ (µ) was 0.65 ± 0.1 and the standard deviation (σ) was 0.17 ± 0.1 (n = 168), consistent with structural predictions for translation-competent 80S ribosomes. **H**. Plots of the 40S subunit recruitment efficiency (left), observed 40S subunit association time (middle), and 60S subunit joining efficiency (right) in the indicated conditions. 40S subunit recruitment efficiency was defined as the fraction of 1,000 analyzed ZMWs with at least one stable (> 10 s) 40S subunit association event. The concentration of 40S subunits (5 nM) was optimized to yield a single recruitment event per ZMW, which is most probable when < 30% of ZMWs contain a recruitment event, as predicted by Poisson distribution statistics. 40S association time (t_40S_) was defined as above (n = 135, 128, 131, 155, and 137, from left to right), and the median observed times are represented by the orange lines. 60S subunit joining efficiency was defined the fraction of recruited 40S subunits (300 analyzed) with a 60S subunit joining event (indicated by Cy3-Cy5 FRET), which was normalized to account for relative labeling efficiencies. **I**. Cumulative probability plot of 40S and 60S subunit association times and the fits to single-exponential functions, which yielded apparent association rates of k_on,40S_ ≈ 0.049 ± 0.002 s^-1^ and k_on,60S_ ≈ 0.033 ± 0.001 s^-1^ when added at 5 nM and 100 nM (final concentration), respectively (n = 128).

**Supplementary Figure 2.**
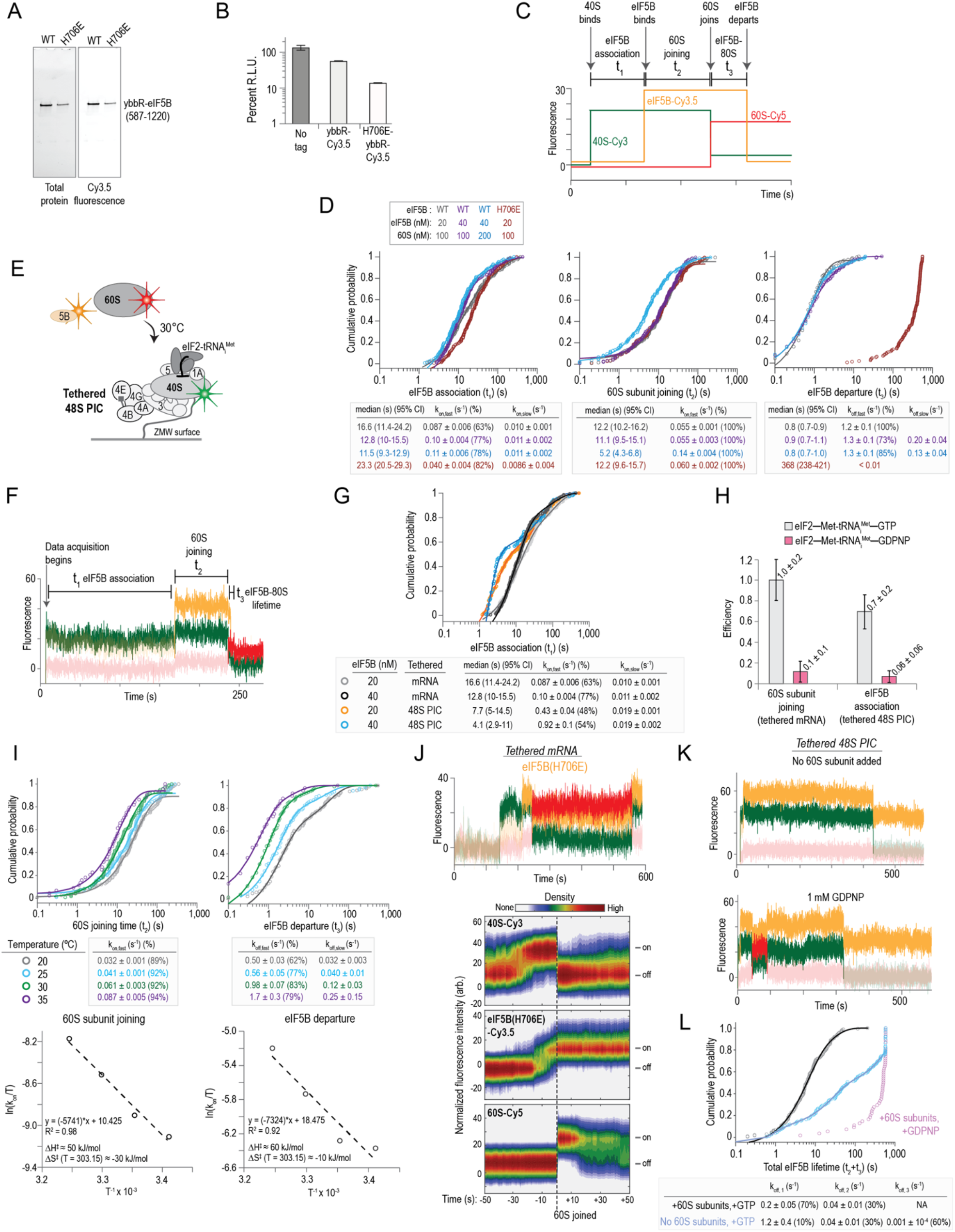
Real-time analysis of eIF5B-mediated ribosomal subunit joining in humans. **A**. Fluorescence scans of gels that analyzed purified and fluorescently-labeled eIF5B proteins. ‘WT’ corresponds to the wild-type protein with the N-terminal domain truncated (residues 1-586) and an N-terminal ybbR peptide tag (11 amino acids). ‘H706E’ corresponds to the GTPase deficient version of the protein. **B**. Plot of the relative light units (RLU) from nanoLuciferase in vitro translation assays where the indicated eIF5B proteins were supplemented in the extract (840 nM final concentration). **C**. Theoretical real-time single-molecule fluorescence data from a ZMW with sequential association of the 40S subunit (green), eIF5B (orange), and 60S subunit (red), which was followed by departure of eIF5B from the newly-formed 80S initiation complex. The dwell times were defined as: between appearance of 40S and eIF5B signals, as the eIF5B association time (t_1_); between appearance of eIF5B and 60S signal, indicated by 40S-60S FRET, as 60S joining time (t_2_); and between appearance of 60S until loss of eIF5B, as eIF5B departure time (t_3_). **D**. Cumulative probability plots of the observed times at the indicated concentrations for eIF5B and 60S subunits at 30 °C. In all experiments, the 43S PIC was present at 10 nM (final concentration, via 40S-Cy3). WT and H706E indicate whether wild-type or catalytically-inactive eIF5B-Cy3.5 were present, respectively. Lines represent fits of observed data to single- or double-exponential functions, which yielded the indicated association or dissociation rates. All errors represent 95% confidence intervals (C.I.). The number of molecules analyzed were: 136 (gray), 148 (purple), 134 (blue), and 146 (red). **E**. Schematic of an alternative experimental setup (‘tethered 48S PIC’) used in some instances to provide sharper focus on eIF5B dynamics and 60S subunit joining. In this setup, a pre-formed 48S PIC at equilibrium on the β-globin mRNA was tethered to the ZMW imaging surface in the presence of 1 mM ATP and GTP. After removal of untethered components, data acquisition began via excitation with a 532 nm laser, and (final concentrations) 20 nM eIF5B-Cy3.5, 100 nM 60S-Cy5 subunits, 1 µM eIF1A, and 1 mM ATP and GTP were added. ZMWs with tethered 48S PICs were identified by the initial presence of 40S-Cy3 fluorescence signal (green). **F**. Example single-molecule fluorescence data from a tethered 48S PIC experiment where eIF5B association led to 60S subunit joining and rapid departure of eIF5B from the 80S initiation complex at 30 °C. The dwells were defined as above in the ‘tethered mRNA’ experimental setup. **G**. Comparison of observed eIF5B association times (t_1_) in tethered mRNA versus tethered 48S PIC experimental setups at 30 °C. The pre-equilibration of the 48S PIC prior to tethering allowed the slow upstream step to proceed, which enabled rapid and concentration-dependent eIF5B association. Fits of the observed association times are represented by the lines, which yielded the indicated rates (n = 172 and 111 for 20 & 40 nM, respectively). The ‘tethered mRNA’ data were replotted from panel B to facilitate comparisons. All errors represent 95% C.I.. **H**. Plot of 60S subunit joining and eIF5B association efficiencies when eIF2–Met-tRNA_iMet_–GTP or eIF2–Met-tRNAiMet–GDPNP were present at 30 °C. 60S subunit joining efficiency was quantified in the tethered mRNA experimental setup with 1 µM unlabeled eIF5B present. eIF5B association efficiency was quantified in the tethered 48S PIC setup (to facilitate analyses) in the absence of 60S subunits. The observed efficiencies and 95% C.I. are indicated. **I**. Cumulative probability (top) and Eyring (bottom) plots of observed 60S joining and eIF5B departure times at 20, 25, 30, and 35 °C. eIF5B-Cy3.5 and 60S-Cy5 subunits were present at 20 nM and 100 nM, respectively. Lines on the cumulative probability plots represent fits of the observed data to single or double-exponential functions, which yielded the indicated rates (errors represent 95% C.I.). Eyring plots (bottom) were modeled via linear regression analyses, which yielded the indicated enthalpies and entropies of activation. The number of molecules analyzed were: 116 (20 °C), 109 (25 °C), 120 (30 °C), and 44 (35 °C). **J**. Example single-molecule fluorescence data (top) and density plots (bottom) from the tethered mRNA setup when catalytically-inactive eIF5B (H706E) was examined at 30 °C. 40S-Cy3, eIF5B(H706E)-Cy3.5, and 60S-Cy5 subunits were present at 10 nM, 20 nM, and 100 nM, respectively. eIF5B(H706E) was unable to depart the 80S initiation complex in the presence of 1 mM GTP. **K**. Example single-molecule fluorescence data from the tethered 48S PIC experimental setup where wild-type eIF5B-Cy3.5 (20 nM) was added to pre-equilibrated 48S PICs either in the absence of 60S subunits (top) or when 60S subunits (100 nM) were present but GTP was replaced with 1 mM GDPNP in free solution (bottom). **L**. Plot of the total lifetime of wild-type eIF5B-Cy3.5 on tethered 48S PICs in the indicated conditions. In the presence of 60S subunits and 1 mM GTP, eIF5B remained on initiation complexes for a total of ∼15 s (t_2_ + t_3_). The total lifetime of eIF5B was lengthened approximately 34-fold to ∼510 s when GTP was replaced with 1 mM of the non-hydrolyzable analog, GDPNP. eIF5B total lifetime also was lengthened dramatically (to ∼ 130 s) when 60S subunits were omitted and 1 mM GTP was present.

**Supplementary Figure 3.**
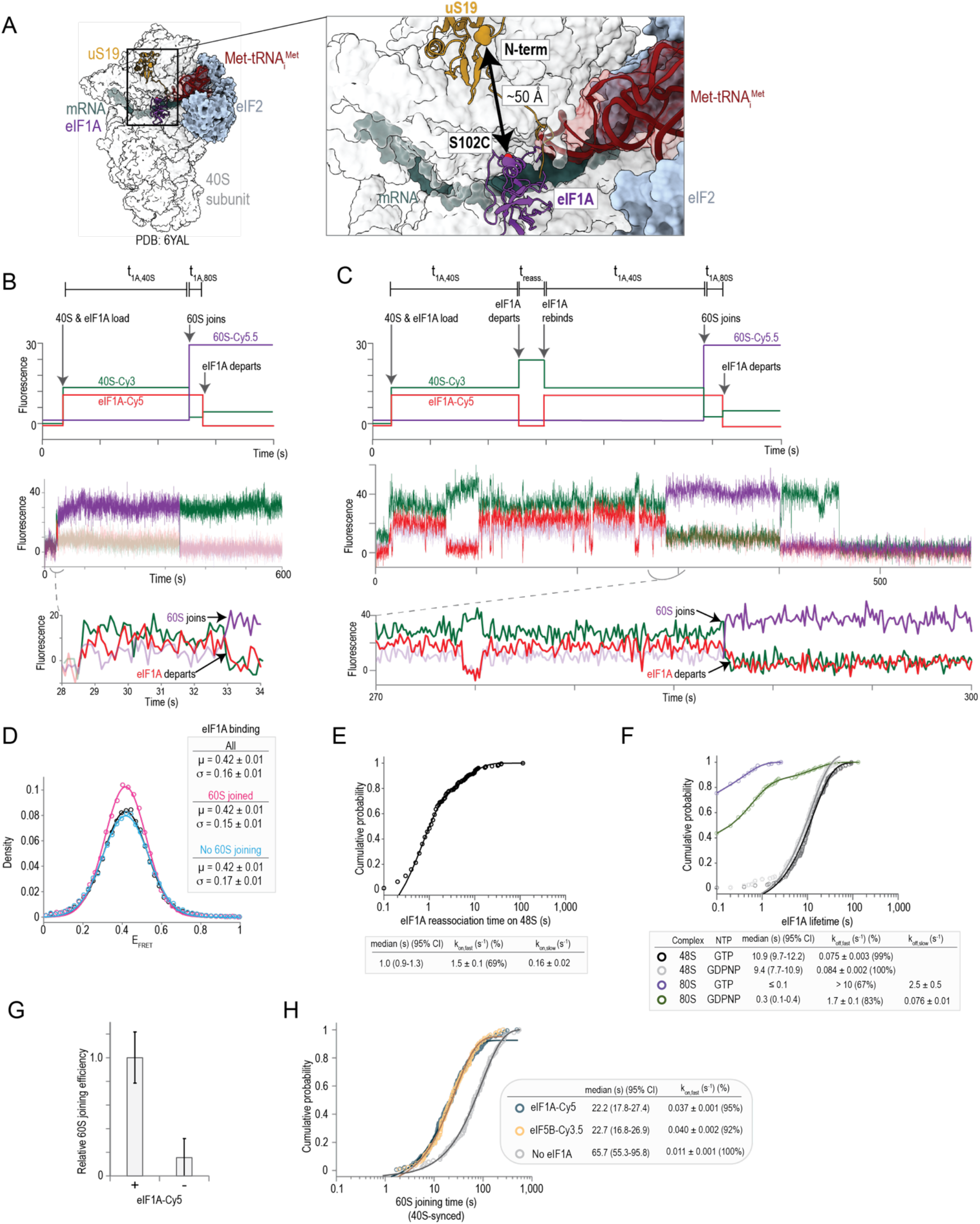
eIF1A resides on initiation complexes until the 60S subunit joins. **A**. Structural model that depicts the proximity of eIF1A and 40S subunit labeling sites. eIF1A was fluorescently labeled via an S102C substitution followed by reaction with Cy5-maleimide dye, which yields a fully-active protein in translation assays, as described previously. 40S subunits were labeled with Cy3 on the N-terminus of uS19, via a fused ybbR tag (11 amino acid), as described previously and used above. The relative proximity of the two labeling sites suggested eIF1A presence on the 40S subunit could be monitored via an unambiguous FRET signal. **B,C**. Theoretical (top) and example (middle, bottom) real-time single-molecule fluorescence data from a ZMW where doubly labeled 43S PICs (10 nM via 40S-Cy3; eIF1A-Cy5), 1 µM unlabeled eIF5B, and 200 nM 60S-Cy5.5 subunits were added to tethered eIF4ABGE-mRNA complexes at 30 °C. During imaging, eIF1A-Cy5 was present at 4.5-fold molar excess relative to 40S subunits. Key association and dissociation events are labeled on the theoretical plot. The lifetime of eIF1A on the 48S PIC (t_1A,48S_) was defined as the duration of individual 40S(Cy3,donor)-eIF1A(Cy5,acceptor) FRET events. In these experiments, 60S subunit joining was indicated by appearance of 40S(Cy3,donor)-60S(Cy5.5,acceptor) FRET. eIF1A lifetime on the 80S complex (t_1A,80S_) was defined as the time elapsed from 60S subunit joining until eIF1A departure (loss of 40S-eIF1A FRET). Panel B depicts an mRNA where eIF1A was co-recruited with the 40S subunit (signal begins in 40S-eIF1A FRET state) and that initial, co-recruited eIF1A protein was present until the 60S subunit joined. Panel C depicts a complex where multiple eIF1A binding events occurred prior to 60S subunit joining. In both cases, eIF1A departed either concomitantly (left) with or within a few hundred milliseconds (right) after 60S subunit joining. **D**. Plot of FRET efficiency (E_FRET_) distributions for 40S(Cy3,donor)-eIF1A(Cy5,acceptor) FRET. ‘All’ represents all observed 40S-eIF1A FRET events (n = 433), ‘60S joined’ represents events that overlapped with 60S subunit joining (n = 175), and ‘No 60S joining’ represents events that did not overlap with 60S subunit joining (n = 258). The lines represent fits of the observed data to single gaussian functions, which yielded the indicated means (µ) and standard deviations (σ). **E**. Cumulative probability plot of observed eIF1A reassociation times with the 48S PIC prior to 60S subunit joining at 30 °C (n = 258). eIF1A-Cy5 was present at ∼45 nM in free solution. The line represents a fit of the observed data to a double-exponential function, which yielded the indicated rates. All errors represent 95% C.I.. **F**. Cumulative probability plot of observed eIF1A lifetimes on either the 48S or 80S initiation complexes when either eIF5B-GTP or eIF5B-GDPNP was present at 30 °C. The line represents a fit of the observed data to either single- or double-exponential functions, which yielded the indicated rates. All errors represent 95% C.I.. **G**. Plot of relative efficiency of 60S subunit joining when eIF1A-Cy5 was present in or eIF1A was absent from initiation reactions. Error bars represent propagated 95% C.I.. **H**. Cumulative probability plot of observed 60S subunit joining times at 30 °C when either 40 nM eIF5B-Cy3.5 (gold, n = 134) or 45 nM eIF1A-Cy5 (slate, n = 175) was present. These findings further confirmed that fluorescently-labeled eIF1A and eIF5B are fully active, as the proteins yielded identical rates of 60S subunit joining as to when the unlabeled version is present (unlabeled eIF1A was present in the eIF5B-Cy3.5 experiment, and vice versa). Moreover, the rare instances where the 60S subunit joined when eIF1A was absent from the reaction occurred 4-fold slower (grey, n = 100). The lines represent fits of the observed data to single-exponential functions, which yielded the indicated rates. All errors represent 95% C.I..

**Supplementary Figure 4.**
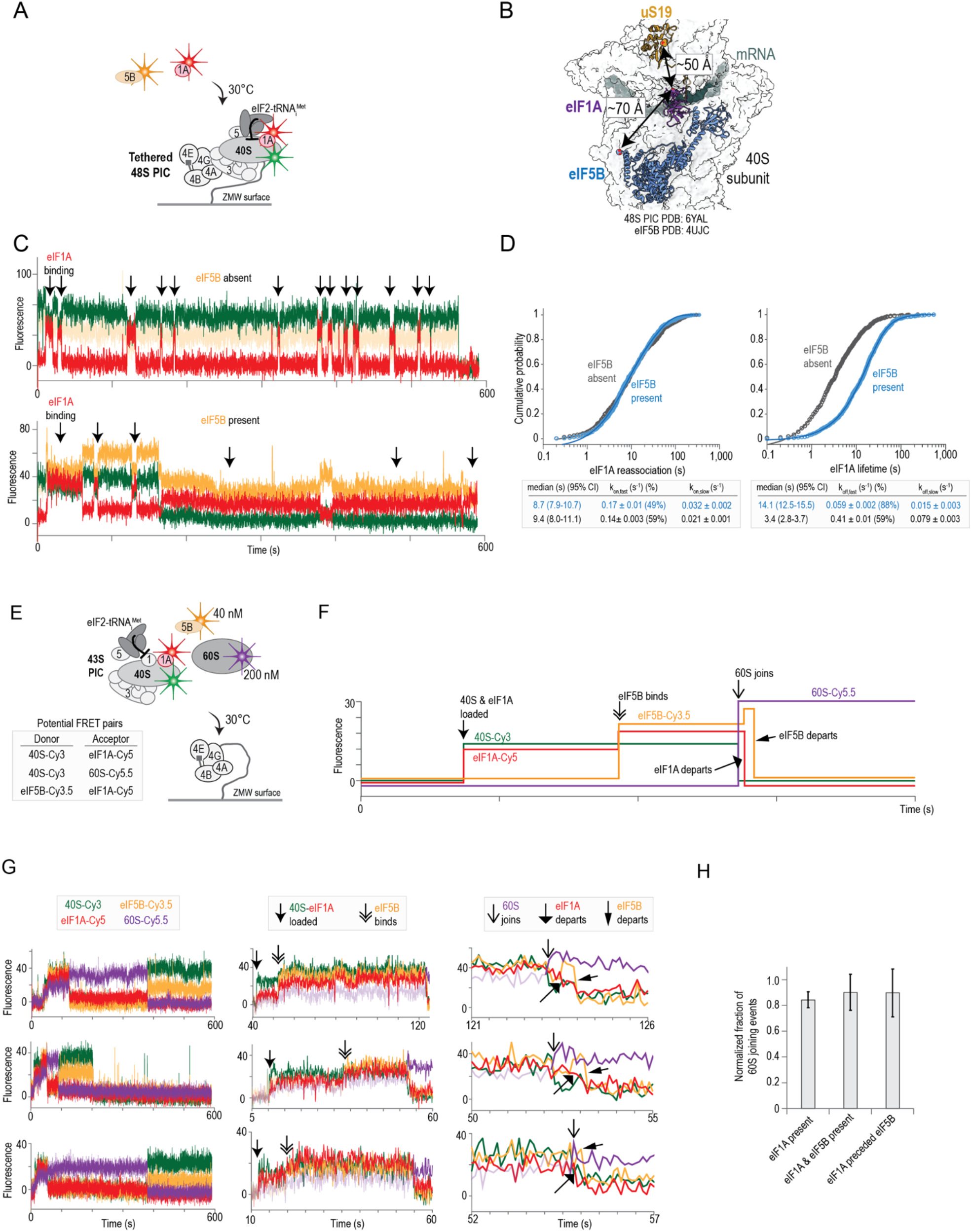
eIF1A and eIF5B reside simultaneously on initiation complexes when the 60S subunit joins. **A**. Schematic of single-molecule experiments that examined eIF1A and eIF5B dynamics on the 48S PIC at equilibrium. Preassembled 48S PICs on β-globin mRNA were tethered at equilibrium within ZMWs in the presence of 1 mM ATP and GTP. After removal of untethered components, data acquisition began via excitation with a 532 nm laser, and an imaging mix that contained (final concentrations) 10 nM eIF1A-Cy5 (red) and 20 nM eIF5B-Cy3.5 (orange) was present at 30 °C. ZMWs with tethered 48S PICs were identified by the initial presence of 40S-Cy3 fluorescence signal (green). **B**. Rudimentary structural comparison where a low-resolution model of human eIF5B (PDB: 4UJC) was docked onto a high-resolution model of a mammalian 48S PIC post-recognition of the start codon (PDB: 6YAL). From these crude analyses, eIF1A is predicted to be within FRET distance (< 80 Å) of the N-terminus of truncated eIF5B, consistent with the eIF5B(Cy3.5, donor)-eIF1A(Cy5,acceptor) FRET signal observed in our single-molecule assays. **C**. Example single-molecule fluorescence data that depicts either: top, a complex with 40S(Cy3,donor)-eIF1A(Cy5,acceptor) FRET in the absence of eIF5B signal, ‘eIF5B absent’; bottom, a complex with both 40S(Cy3,donor)-eIF1A(Cy5,acceptor) and eIF5B(Cy3.5,donor)-eIF1A(Cy5,acceptor) FRET, ‘eIF5B present’. Focused analyses were conducted on both forms of the 48S PIC to derive kinetic parameters. **D**. Cumulative probability plots of observed eIF1A reassociation times with (left) or eIF1A lifetimes on (right) the 48S PIC at 30 °C at equilibrium. eIF1A-Cy5 was present at 10 nM. The lines represent fits of the observed data to double-exponential functions, which yielded the indicated rates. All errors represent 95% C.I.. 831 and 589 eIF1A binding events were analyzed when eIF5B was present or absent, respectively. **E**. Schematic of a four-color single-molecule experiment. The doubly labeled 43S PIC (10 nM by 40S-Cy3; eIF1A-Cy5), 40 nM eIF5B-Cy3.5, and 200 nM 60S-Cy5.5 subunit were added to β-globin mRNA tethered within ZMWs in the presence of saturating concentrations of eIF4ABGE and 1 mM ATP and GTP at 30 °C. During imaging, eIF1A-Cy5 was present at 4.5-fold molar excess relative to the 40S subunit. Fluorescence data were acquired for 600 s with excitation via the 532 nm laser. The potential FRET signals are indicated in the box. **F,G**. Theoretical (panel F) and example (panel G) four-color single-molecule fluorescence data from a ZMW where a recruited doubly labeled 43S PIC (40S-Cy3, green; eIF1A-Cy5, red) was bound by eIF5B-Cy3.5 (orange), which was followed by 60S-Cy5.5 (purple) subunit joining. The 43S PIC was recruited in a 40S(Cy3, donor)-eIF1A(Cy5, acceptor) FRET state. Once eIF5B bound, the eIF1A-Cy5 signal increased due to eIF5B(Cy3.5, donor)-eIF1A(Cy5, acceptor) FRET. Joining of the 60S subunit was indicated by appearance of Cy5.5 fluorescence signal due to 40S(Cy3, donor)-60S(Cy5.5, acceptor) FRET. Departure of eIF1A-Cy5 and eIF5B-Cy3.5 was indicated by loss of Cy5 and Cy3.5 fluorescence signals, respectively. In panel G, the full experimental window is depicted on the left, and the middle and right panels represent zoomed views of the indicated time windows. Given bleed through across the four fluorescent channels, the fluorescence signals in each channel were made transparent before relevant events for presentation here. **H**. Quantification of eIF1A and eIF5B occupancy upon 60S subunit joining in the four-color single-molecule experiment. The potentially convoluted FRET signals and fluorescence bleed through among channels and factors precluded rigorous kinetic analyses, as exact frames for association and dissociation were extremely challenging to assign. However, the experiment did allow the presence of eIF1A and eIF5B upon 60S subunit joining to be quantified unmistakably. On a large majority of 48S initiation complexes (183/271), eIF1A-Cy5 signal was present when the 60S subunit joined, which corresponded to an estimated eIF1A occupancy of about 85 ± 5 % after correction for eIF1A labeling efficiency (70-90%). Of those eIF1A-bound 48S complexes, nearly half (88/183) also contained eIF5B-Cy3.5 signal when the 60S subunit joined, which indicated that about 90 ± 10 % of the 48S PICs contained both eIF1A and eIF5B, after correction for eIF5B labeling efficiency (∼45%). eIF1A preceded eIF5B association on nearly all (79/88; 90 ± 20 %) 48S PICs that contained both labeled protein when the 60S subunit joined.

**Supplementary Figure 5.**
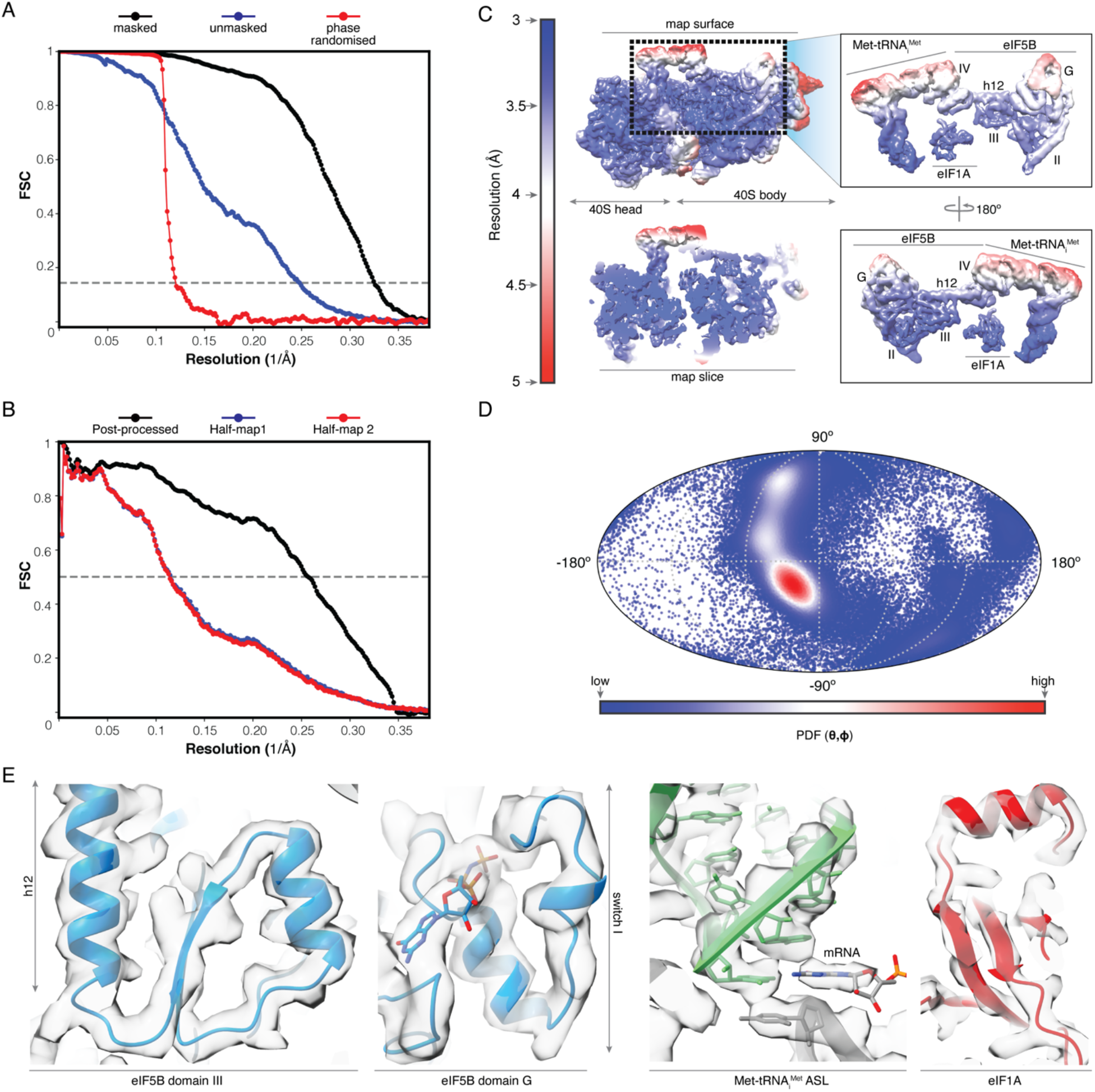
Fourier Shell Correlation curves, local resolution, particle distribution plot and representative cryo-EM density examples. **A**. Fourier Shell Correlation (FSC) curves computed for independently refined half-maps for the final subgroup of particles before masking (blue) and after masking (black). The red curve corresponds to FSC curve of phase-randomized structure factors beyond 7Å. **B**. Model versus map FSC (black). Red and blue curves correspond to a model-map overfitting validation test performed for the final model. A per-atom, random distortion of 0.5Å was introduced in the model which was subsequentially refined against half-map 1 only. FSC between the distorted and refined model against half-map 1 (blue) and half-map 2 (red, not included in the refinement) are nearly identical what guarantees absence of overfitting in the final model. **C**. Unsharpened map colored according to local resolution calculations with the resolution scale indicated on the left and a zoomed view for Met-tRNAiMet/eIF1A/eIF5B on the right. **D**. Eulerian angular distribution for the final refined set of particles in a mollweide spherical projection. **E**. Several views of the final post-processed cryo-EM density used for model building and refinement. The areas of the map to which the different views belong are indicated.

**Supplementary Figure 6.**
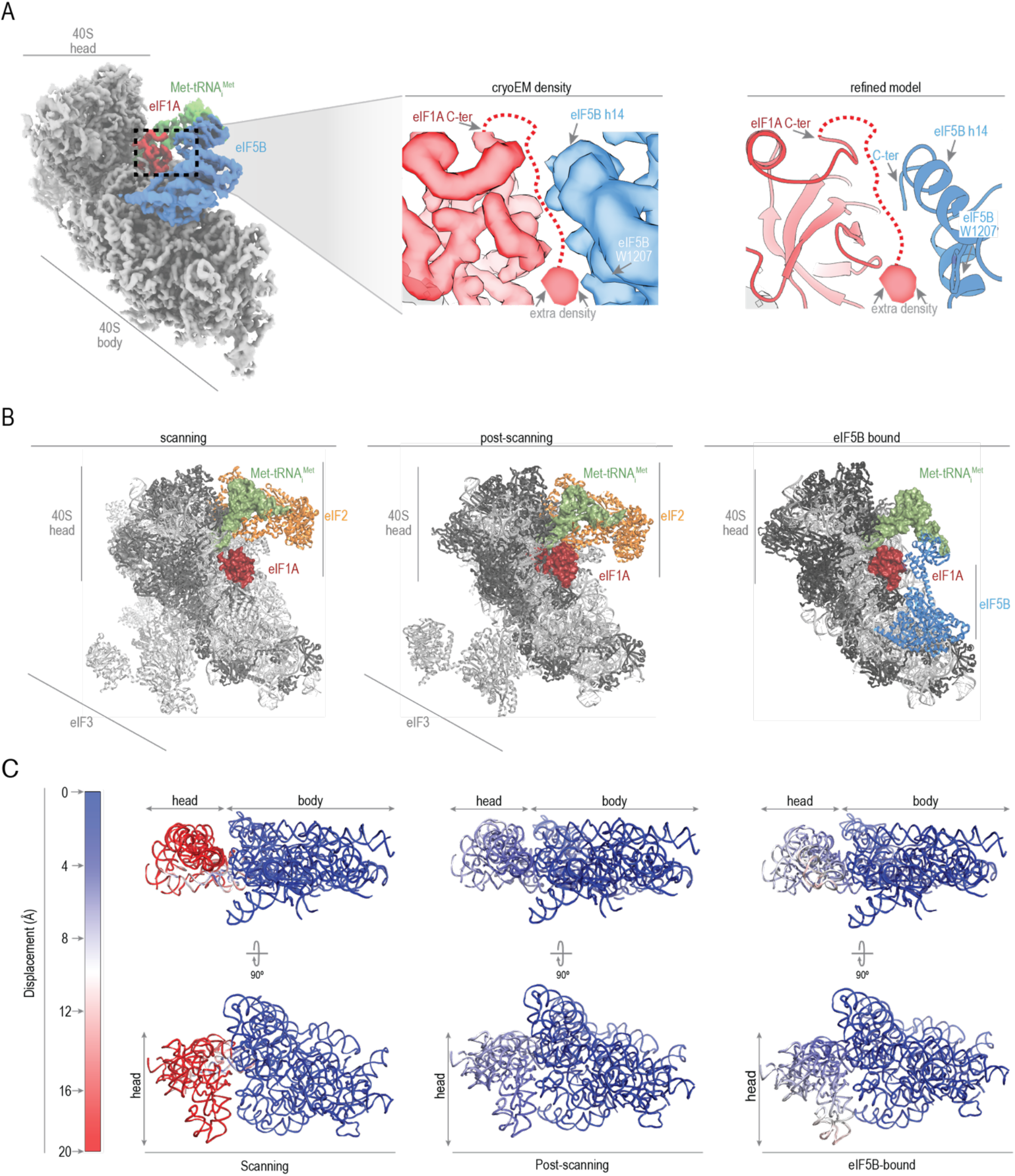
**A**. Post-processed cryo-EM density of eIF1A and eIF5B and the resulting models. While nearly the entire flexible C-terminal tail of eIF1A was disordered, extra density proximal to W1207 of eIF5B was observed, which likely corresponds to I141 in the eIF1A C-terminal tail. The areas of the map to which the different views belong are indicated. **B**. Position of eIF1A and Met-tRNAi^Met^ in scanning, post-scanning, and eIF5B-bound initiation complexes. eIF1A remains bound in the ribosomal A site throughout initiation, until the 60S subunit joins. **C**. Structural models of the 40S subunit (with ribosomal proteins omitted) that indicate movement of the 18S rRNA in the indicated states relative to the final position in the elongation-competent 80S ribosome. In both the post-scanning and eIF5B-bound states, the 40S head region resembles the conformation in the 80S ribosome.

**Supplementary Figure 7.**
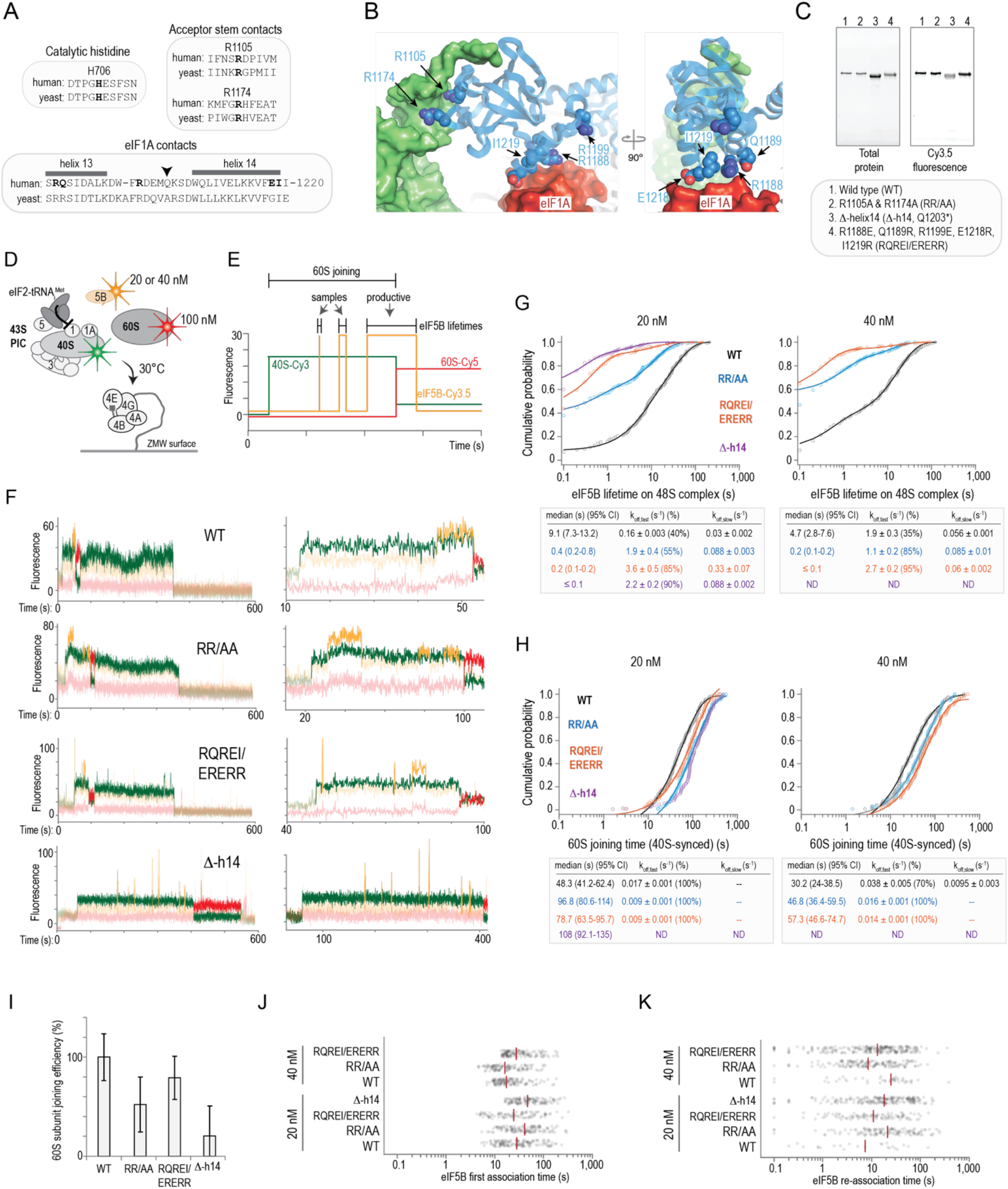
eIF5B contacts with eIF1A and Met-tRNA_i_^Met^ mediate 60S subunit joining. **A**. Alignments of human and yeast eIF5B protein sequences at the indicated regions. **B**. Views of the eIF5B-bound 48S complex that depict the location of R1105 and R1174, which contact Met-tRNAiMet, and R1188, Q1189, R1199, E1218, and I1219, which contact eIF1A. **C**. Fluorescence scans of a gel that analyzed the indicated purified and fluorescently-labeled eIF5B proteins. All proteins had the N-terminal domain removed (residues 1-586) and contained an N-terminal ybbR peptide tag (11 amino acids). **D**. Schematic of the single-molecule experiment to assess the role of the indicated residues in eIF5B. The 43S PIC (10 nM via 40S-Cy3 subunits, green), wild-type or mutant eIF5B-Cy3.5 (orange, at 20 or 40 nM), and 100 nM 60S-Cy5 (red) subunit were added to β-globin mRNA tethered within ZMWs in the presence of saturating concentrations of eIF4ABGE and 1 mM ATP and GTP at 30 °C. Fluorescence data were acquired for 600 s with excitation via the 532 nm laser. **E**. Theoretical real-time single-molecule fluorescence data from a ZMW with sequential association of the 40S subunit (green), eIF5B (orange), and 60S subunit (red), which was followed by departure of eIF5B from the newly-formed 80S initiation complex. In this experiment, the lifetimes of all eIF5B binding events that began prior to 60S subunit joining were analyzed. Events that concluded prior to 60S subunit joining were classified as ‘samples’, and events that concluded after 60S subunit joining were classified as ‘productive’. To facilitate cross-comparisons in these experiments, the 60S subunit joining time was defined as the time elapsed from 40S subunit loading (appearance of green fluorescence) until 60S subunit joining, as indicated by 40S(Cy3, donor)-to-60S(Cy5, acceptor) FRET. **F**. Example single-molecule fluorescence data obtained when the indicated eIF5B proteins were present. The left plots represent the fluorescence intensities throughout the entire 600 s data collection window, and the right are zoomed views of events of interest. **G,H**. Cumulative probability plots of observed eIF5B lifetimes on the 48S complex (panel G) or the 60S subunit joining time (relative to 40S subunit loading) (panel H) when the indicated eIF5B protein was present at the indicated concentration (20 or 40 nM). The lines represent fits of the observed data to double-exponential functions, which yielded the indicated rates. All errors represent 95% C.I.. **I**. Plot of the 60S subunit joining efficiency in ZMWs that contained at least one 40S subunit loading event when the indicated eIF5B proteins were present at 40 nM. **J,K**. Plots of the observed times for the first eIF5B binding event to occur (relative to 40S subunit loading) (panel J) and subsequent eIF5B binding events to occur (relative to the previous eIF5B binding event) (panel K) when the indicated eIF5B proteins were present at 20 or 40 nM. Medians are indicated by the orange line.

**Supplementary Table S1.**
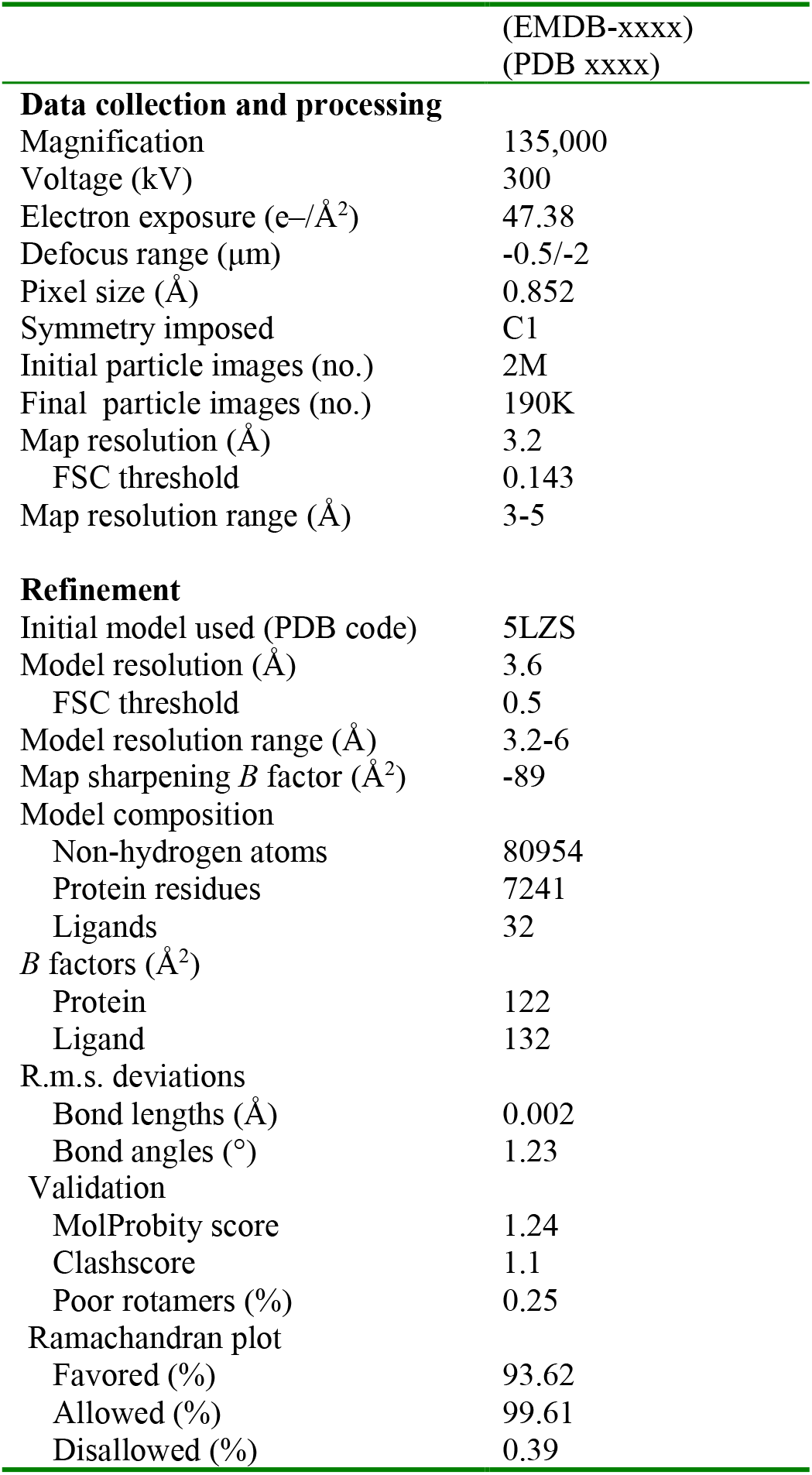
Cryo-EM data collection, refinement and validation statistics.

**Supplementary Movie 1**.

Proposed model for how human eIF1A and eIF5B collaborate to orient Met-tRNA_i_^Met^ properly and catalyze ribosomal subunit joining, as illuminated by our single-molecule dynamics and structural results. The 40S LIC presented here was compared with a previous yeast eIF5B-bound 80S initiation complex *(13)*.

## REFERENCES

1. V. M. Pain, Initiation of Protein Synthesis in Eukaryotic Cells. European Journal of Biochemistry 236, 747–771 (1996).

2. W. C. Merrick, G. D. Pavitt, Protein Synthesis Initiation in Eukaryotic Cells. Cold Spring Harb Perspect Biol 10, (2018).

3. J. W. B. Hershey, N. Sonenberg, M. B. Mathews, Principles of Translational Control. Cold Spring Harbor perspectives in biology 11, (2019).

4. C. E. Aitken, J. R. Lorsch, A mechanistic overview of translation initiation in eukaryotes. Nature Structural and Molecular Biology 19, 568–576 (2012).

5. M. Sokabe, C. S. Fraser, Toward a Kinetic Understanding of Eukaryotic Translation. Cold Spring Harbor perspectives in biology 11, (2019).

6. A.G. Hinnebusch, The scanning mechanism of eukaryotic translation initiation. Annual Review of Biochemistry 83, 779–812 (2014).

7. A. G. Hinnebusch, Structural Insights into the Mechanism of Scanning and Start Codon Recognition in Eukaryotic Translation Initiation. Trends in biochemical sciences 42, 589–611 (2017).

8. Y. Hashem, J. Frank, The Jigsaw Puzzle of mRNA Translation Initiation in Eukaryotes: A Decade of Structures Unraveling the Mechanics of the Process. Annual review of biophysics 47, 125–151 (2018).

9. T. V. Pestova et al., The joining of ribosomal subunits in eukaryotes requires eIF5B. Nature 403, 332–335 (2000).

10. J. H. Lee et al., Initiation factor eIF5B catalyzes second GTP-dependent step in eukaryotic translation initiation. Proc Natl Acad Sci U S A 99, 16689–16694 (2002).

11. A. S. Shin et al., Uncoupling of initiation factor eIF5B/IF2 GTPase and translational activities by mutations that lower ribosome affinity. Cell 111, 1015–1025 (2002).

12. I.S. Fernández et al., Molecular architecture of a eukaryotic translational initiation complex. Science 342, 1240585 (2013).

13. Wang et al., Structural basis for the transition from translation initiation to elongation by an 80S-eIF5B complex. Nature Communications 11, (2020).

14. H. Yamamoto et al., Structure of the mammalian 80S initiation complex with initiation factor 5B on HCV-IRES RNA. Nature Structural and Molecular Biology 21, 721–727 (2014).

15. S. K. Choi et al., Physical and Functional Interaction between the Eukaryotic Orthologs of Prokaryotic Translation Initiation Factors IF1 and IF2. Molecular and Cellular Biology 20, 7183–7191 (2000).

16. S. Olsen, Domains of eIF1A that mediate binding to eIF2, eIF3 and eIF5B and promote ternary complex recruitment in vivo. The EMBO Journal 22, 193–204 (2003).

17. M. G. Acker, B.-S. Shin, T. E. Dever, J. R. Lorsch, Interaction between Eukaryotic Initiation Factors 1A and 5B Is Required for Efficient Ribosomal Subunit Joining. Journal of Biological Chemistry 281, 8469–8475 (2006).

18. M. Fringer, M. G. Acker, C. A. Fekete, J. R. Lorsch, T. E. Dever, Coupled Release of Eukaryotic Translation Initiation Factors 5B and 1A from 80S Ribosomes following Subunit Joining. Molecular and Cellular Biology 27, 2384–2397 (2007).

19. V. P. Pisareva, A. V. Pisarev, eIF5 and eIF5B together stimulate 48S initiation complex formation during ribosomal scanning. Nucleic Acids Research 42, 12052–12069 (2014).

20. J. Chen et al., High-throughput platform for real-time monitoring of biological processes by multicolor single-molecule fluorescence. Proceedings of the National Academy of Sciences 111, 664–669 (2014).

21. C. P. Lapointe et al., Dynamic competition between SARS-CoV-2 NSP1 and mRNA on the human ribosome inhibits translation initiation. Proceedings of the National Academy of Sciences 118, e2017715118 (2021).

22. A. Koch, L. Aguilera, T. Morisaki, B. Munsky, T. J. Stasevich, Quantifying the dynamics of IRES and cap translation with single-molecule resolution in live cells. Nat Struct Mol Biol 27, 1095–1104 (2020).

23. J. Wang et al., eIF5B gates the transition from translation initiation to elongation. Nature 573, 605–608 (2019).

24. Sokabe, C. S. Fraser, Human eukaryotic initiation factor 2 (eIF2)-GTP-Met-tRNAi ternary complex and eIF3 Stabilize the 43 S preinitiation complex. Journal of Biological Chemistry 289, 31827–31836 (2014).

25. G. Acker et al., Kinetic Analysis of Late Steps of Eukaryotic Translation Initiation. Journal of Molecular Biology 385, 491–506 (2009).

26. L. A. Passmore et al., The eukaryotic translation initiation factors eIF1 and eIF1A induce an open conformation of the 40S ribosome. Molecular cell 26, 41–50 (2007).

27. Y. Hashem et al., Structure of the mammalian ribosomal 43S preinitiation complex bound to the scanning factor DHX29. Cell 153, 1108–1119 (2013).

28. I. B. Lomakin, T. A. Steitz, The initiation of mammalian protein synthesis and mRNA scanning mechanism. Nature 500, 307–311 (2013).

29. Weisser, F. Voigts-Hoffmann, J. Rabl, M. Leibundgut, N. Ban, The crystal structure of the eukaryotic 40S ribosomal subunit in complex with eIF1 and eIF1A. Nature Structural & Molecular Biology 20, 1015–1017 (2013).

30. T. Hussain et al., Structural changes enable start codon recognition by the eukaryotic translation initiation complex. Cell 159, 597–607 (2014).

31. A.Des Georges et al., Structure of mammalian eIF3 in the context of the 43S preinitiation complex. Nature 525, 491–495 (2015).

32. A. Simonetti et al., eIF3 Peripheral Subunits Rearrangement after mRNA Binding and Start-Codon Recognition. Mol Cell 63, 206–217 (2016).

33. J.L. Llácer et al., Translational initiation factor eIF5 replaces eIF1 on the 40S ribosomal subunit to promote start-codon recognition. eLife 7, 1–33 (2018).

34. J. Brito Querido et al., Structure of a human 48S translational initiation complex. Science (New York, N.Y.) 369, 1220–1227 (2020).

35. A. Simonetti, E. Guca, A. Bochler, L. Kuhn, Y. Hashem, Structural Insights into the Mammalian Late-Stage Initiation Complexes. Cell reports 31, 107497–107497 (2020).

36. H. Kratzat et al., A structural inventory of native ribosomal ABCE1-43S pre-initiation complexes. EMBO J 40, e105179 (2021).

37. N. Nag et al., eIF1A/eIF5B interaction network and its functions in translation initiation complex assembly and remodeling. Nucleic Acids Research, gkw552 (2016).

38. A. Marintchev, V. G. Kolupaeva, T. V. Pestova, G. Wagner, Mapping the binding interface between human eukaryotic initiation factors 1A and 5B: A new interaction between old partners. Proceedings of the National Academy of Sciences 100, 1535–1540 (2003).

39. S. Shao et al., Decoding Mammalian Ribosome-mRNA States by Translational GTPase Complexes. Cell 167, 1229-1240.e1215 (2016).

40. A. V. Pisarev et al., Specific functional interactions of nucleotides at key -3 and +4 positions flanking the initiation codon with components of the mammalian 48S translation initiation complex. Genes Dev 20, 624–636 (2006).

41. S. Suresh et al., eIF5B drives integrated stress response-dependent translation of PD-L1 in lung cancer. Nature cancer 1, 533–545 (2020).

42. A. Roll-Mecak, C. Cao, T. E. Dever, S. K. Burley, X-Ray Structures of the Universal Translation Initiation Factor IF2/eIF5B. Cell 103, 781–792 (2000).

43. B. Kuhle, R. Ficner, eIF5B employs a novel domain release mechanism to catalyze ribosomal subunit joining. The EMBO Journal 33, 1177–1191 (2014).

44. A. Zheng et al., X-ray structures of eIF5B and the eIF5B–eIF1A complex: the conformational flexibility of eIF5B is restricted on the ribosome by interaction with eIF1A. Acta Crystallographica Section D Biological Crystallography 70, 3090–3098 (2014).

45. S. Kaledhonkar et al., Late steps in bacterial translation initiation visualized using time-resolved cryo-EM. Nature 570, 400–404 (2019).

46. R. A. Marshall, C. E. Aitken, J. D. Puglisi, GTP Hydrolysis by IF2 Guides Progression of the Ribosome into Elongation. Molecular Cell 35, 37–47 (2009).

47. C. Ling, D. N. Ermolenko, Initiation factor 2 stabilizes the ribosome in a semirotated conformation. Proceedings of the National Academy of Sciences 112, 15874–15879 (2015).

48. T. Sprink et al., Structures of ribosome-bound initiation factor 2 reveal the mechanism of subunit association. Sci Adv 2, e1501502 (2016).

49. A. G. Johnson et al., RACK1 on and off the ribosome. RNA, rna.071217.071119-rna.071217.071119 (2019).

50. J. Yin, A. J. Lin, D. E. Golan, C. T. Walsh, Site-specific protein labeling by Sfp phosphopantetheinyl transferase. Nature Protocols 1, 280–285 (2006).

51. C. S. Fraser, K. E. Berry, J. W. B. Hershey, J. A. Doudna, eIF3j Is Located in the Decoding Center of the Human 40S Ribosomal Subunit. Molecular Cell 26, 811–819 (2007).

52. A.R. Özeş, K. Feoktistova, B. C. Avanzino, C. S. Fraser, Duplex unwinding and ATPase activities of the DEAD-box helicase eIF4A are coupled by eIF4G and eIF4B. J Mol Biol 412, 674–687 (2011).

53. Feoktistova, E. Tuvshintogs, A. Do, C. S. Fraser, Human eIF4E promotes mRNA restructuring by stimulating eIF4A helicase activity. Proc Natl Acad Sci U S A 110, 13339–13344 (2013).

54. Sokabe, C. S. Fraser, J. W. Hershey, The human translation initiation multi-factor complex promotes methionyl-tRNAi binding to the 40S ribosomal subunit. Nucleic Acids Res 40, 905–913 (2012).

55. Damoc et al., Structural characterization of the human eukaryotic initiation factor 3 protein complex by mass spectrometry. Molecular & cellular proteomics : MCP 6, 1135–1146 (2007).

56. J. Chen et al., High-throughput platform for real-time monitoring of biological processes by multicolor single-molecule fluorescence. Proceedings of the National Academy of Sciences of the United States of America 111, 664–669 (2014).

57. C. E. Aitken, R. A. Marshall, J. D. Puglisi, An oxygen scavenging system for improvement of dye stability in single-molecule fluorescence experiments. Biophysical Journal 94, 1826–1835 (2008).

58. F. Juette et al., Single-molecule imaging of non-equilibrium molecular ensembles on the millisecond timescale. Nature Methods 13, 341–344 (2016).

59. J. E. Bronson, J. Fei, J. M. Hofman, R. L. Gonzalez, C. H. Wiggins, Learning rates and states from biophysical time series: a Bayesian approach to model selection and single-molecule FRET data. Biophys J 97, 3196–3205 (2009).

60. M. R. Lawson et al., Mechanisms that ensure speed and fidelity in eukaryotic translation termination. Science 373, 876–882 (2021).

61. J. Zivanov, T. Nakane, S. H. W. Scheres, Estimation of high-order aberrations and anisotropic magnification from cryo-EM data sets in. IUCrJ 7, 253–267 (2020).

62. A. Rohou, N. Grigorieff, CTFFIND4: Fast and accurate defocus estimation from electron micrographs. J Struct Biol 192, 216–221 (2015).

63. A. Casañal, B. Lohkamp, P. Emsley, Current developments in Coot for macromolecular model building of Electron Cryo-microscopy and Crystallographic Data. Protein Sci 29, 1069–1078 (2020).

64. J. Jumper et al., Highly accurate protein structure prediction with AlphaFold. Nature 596, 583–589 (2021).

65. V. Afonine et al., Real-space refinement in PHENIX for cryo-EM and crystallography. Acta Crystallogr D Struct Biol 74, 531–544 (2018).

66. R. A. Nicholls, F. Long, G. N. Murshudov, Low-resolution refinement tools in REFMAC5. Acta Crystallogr D Biol Crystallogr 68, 404–417 (2012).

67. J. Williams et al., MolProbity: More and better reference data for improved all-atom structure validation. Protein Sci 27, 293–315 (2018).

68. T. D. Goddard et al., UCSF ChimeraX: Meeting modern challenges in visualization and analysis. Protein science : a publication of the Protein Society 27, 14–25 (2018).

